# Blockade of rheumatoid arthritis synovial fluid-induced sensory neuron activation by JAK inhibitors

**DOI:** 10.1101/2024.08.19.608613

**Authors:** Yuening Li, Elizabeth H. Gray, Rosie Ross, Irene Zebochin, Amy Lock, Laura Fedele, Louisa Janice Kamajaya, Rebecca J. Marrow, Sarah Ryan, Pascal Röderer, Oliver Brüstle, Susan John, Franziska Denk, Leonie S. Taams

## Abstract

**Objective:** Clinical studies suggest that compared to anti-TNF treatment, JAK inhibitors (JAKi) are superior in reducing pain in rheumatoid arthritis (RA). The underlying mechanisms for this observation are still unknown. Sensory neurons transmit noxious signals from inflamed joints to the central nervous system, where a pain percept is generated. We investigated whether JAKi exert direct effects on sensory neurons.

**Methods:** In-house and public RNA sequencing datasets of sensory neurons were analysed for relevant JAK/STAT and cytokine-receptor gene expression. Human induced pluripotent stem cell (IPSC)-derived sensory neurons were stimulated with serum and synovial fluid (SF) from individuals with RA, or with selected cytokines that were found in RA SF by Luminex. Phosphorylation of STAT3 (pSTAT3) was assessed by Western blot. Sensory neuron activation was examined by recording neuronal firing using multi-electrode array and measuring expression levels of pain-relevant genes with STAT3-binding sites.

**Results:** Cell-free RA synovial fluid induced pSTAT3 in IPSC-derived sensory neurons, an effect which was completely blocked by the JAKi tofacitinib. Compared to paired serum, RA SF was enriched for the JAK/STAT cytokines IL-6, IL-11, LIF, IFN-alpha and IFN-beta, with their requisite receptors present on sensory neurons. Stimulation of IPSC-derived sensory neurons with these recombinant cytokines recapitulated pSTAT3 induction in these cells. Furthermore, IL-6+sIL-6R or LIF upregulated expression of pain-relevant genes which was blocked by tofacitinib. Finally, we provided evidence that LIF can induce neuronal sensitisation.

**Conclusion:** Our data indicate that JAKi can act directly on sensory neurons, providing a potential mechanistic explanation for their suggested superior analgesic properties.

## Introduction

Rheumatoid arthritis (RA) is a chronic autoimmune disease characterised by inflammation, stiffness and pain of bilateral joints ^4,5^. The advent of biologics has revolutionised the treatment of RA, leading to substantial reduction in overall disease activity. However, moderate to severe pain persists in many patients despite improved treatments and remains a critical unmet need for individuals living with RA ^6,7^.

Amongst current treatments, a class of small-molecule drugs that inhibit Janus kinase (JAK) proteins, called JAK inhibitors (JAKi), show promising results in terms of pain relief. JAKi work by inhibiting the ATP binding domains of JAK, disabling cytokine binding-induced phosphorylation and its downstream effects ^8^. Multiple cytokines signal through JAK proteins and their partner, Signal Transducer and Activator of Transcription (STAT) molecules. JAKi can block multiple cytokines simultaneously, which may underlie their high efficacy ^9^. Several lines of evidence suggest that JAKi may be superior to other treatments, including anti-TNF, in reducing pain in RA. Head-to-head studies showed that the clinically approved JAKi, upadacitinib and baricitinib, performed better on secondary outcome measures such as worst joint pain and mean visual analogue scale for pain ^10,11^. Post-hoc mediator analyses also suggested that baricitinib may directly inhibit pain, as pain reduction exceeded the reduction in inflammatory markers in individuals with RA ^12^.

The mechanisms behind the proposed superior analgesic efficacy of JAKi remain unclear. Pain in RA is initiated by the activation of sensory neurons that innervate inflamed joints and transmit noxious information to the central nervous system, where the pain percept is generated ^13^. Many mechanistic studies of JAKi have focused on their inhibition of immune cell activation and proliferation ^8,14^, which would indirectly reduce pain. It has also been the suggested that JAKi might be able to act directly on pain pathways in the central nervous system ^15^. Here, we explored the possibility that the same might be true for peripheral sensory neurons - the first responders that become aberrantly activated in pain states. Specifically, we hypothesised that JAK/STAT signalling cytokines in the inflamed RA joint exert direct effects on sensory neuron activation, which can be blocked by JAKi.

We tested this hypothesis using induced pluripotent stem cells (IPSC)-derived sensory neurons as a model system. Our findings demonstrate that JAK signalling is active in sensory neurons, and can be driven by multiple cytokines present in the joint fluid of individuals with RA. We propose this direct action on sensory neurons as one putative explanation for the suggested superior analgesic efficacy of JAKi.

## Materials and Methods

All reagents and antibodies used, including their concentrations and catalogue numbers are listed in **Supplementary Table 1**. Fibroblasts, PBMC/SFMC culture, sensitivity analyses and bioinformatic analyses are described in the **Supplementary Methods.**

### IPSC culture, sensory neuron differentiation and maintenance

Two IPSC lines, Kute4 (HPSI0714i-kute_4, female) and UKB (UKBi013-A-GCamP6f, male ^16^) were used in this study. IPSC were maintained in Stemflex media on vitronectin-coated wells and passaged using versene when reaching ∼70% confluency. IPSC were differentiated into sensory neurons once reaching ∼60% confluency following the Chambers protocol and replated at 75,000 cells per coverslip in a 24-well plate at day 11 ^17,18^. Neurons were cultured in N2 media supplemented with NGF, NT-3, GDNF and BDNF at 25 ng/mL (day 11-39) or 10ng/mL (day 39 onwards). 1-3 μM AraC was supplemented to inhibit the growth of non-neuronal cells based on morphological examination around day 14 to day 30. Geltrex (1/250) was supplemented once every week to stabilise the culture and prevent peeling.

### Human samples

Ethics approval for the study and sample collection was obtained from the Bromley Research Ethics Committee (06/Q0705/20) and Harrow Research Ethics Committee (17/LO/1940). For healthy control samples, following written informed consent, a maximum of 100mL peripheral blood was drawn from healthy donors into Vacutainer tubes (BD Biosciences) or in some instances, leucocyte cones from healthy volunteer blood donations were purchased from the NHS Blood and Transplant service (Oxford, UK). Peripheral blood samples and synovial fluid samples from RA patients were collected with informed consent from patients attending Guy’s Hospital Rheumatology Department (London, UK). RA synovial fibroblasts were derived from joint replacement surgery or synovial tissue biopsies and cryopreserved in liquid nitrogen. Details of PBMC, SFMC and synovial fibroblast isolation and cultures can be found in the **Supplementary Methods**. Cell-free serum was obtained by centrifuging a serum tube at 3000*g* for 10 minutes and collecting the supernatant. Cell-free synovial fluid (SF) was obtained by centrifuging SF at 5900*g* for 3.5 min and extracting the supernatant. Cell-free serum and SF aliquots were stored at −80°C.

### Neuron stimulation

Neurons past differentiation day 50 were used in this study. For SF stimulation, SF was centrifuged at 16000*g* for 4 minutes and diluted to 10% in N2 media. For cytokine stimulation, neurons were incubated in the cytokines for the time and concentration specified in the figure legends. In experiments using JAKi, neurons were pre-incubated with N2 media containing tofacitinib (tofa), baricitinib (bari), or upadacitinib (upa) for 1 hour before cytokine stimulation. IL-6 and sIL-6R, and IL-11 and sIL-11R were pre-mixed for 1 hour prior to use in experiments. The cytokine stimulation media also contained the same concentration of JAKi to ensure continuous blockade.

### Immunocytochemistry

Neuron-containing coverslips were fixed in 4% paraformaldehyde (PFA) at room temperature for 10 minutes. Purity staining (BRN3A and PGP9.5) was performed on neurons blocked with 10% donkey serum in 0.1% PBS-Triton for 1 hour, then incubated with primary antibodies overnight at room temperature (BRN3A, PGP9.5). After washes, secondary antibodies were applied for 2 hours at room temperature. To stain for pSTAT3, BRN3A and NF200, following PFA fixation, cells were further permeabilised in 100% methanol at −20°C for 10 minutes, followed by blocking in 5% donkey serum in 0.3% PBS-Triton for 1 hour and incubation with primary antibodies overnight (pSTAT3, BRN3A, NF200) at 4°C. Secondary antibodies were applied for 2 hours, followed by a final incubation in horse-anti-rabbit streptavidin for 30 minutes. PBS washes were performed between each step. Coverslips were mounted on DAPI containing mounting media. The proportion of BRN3A+ neurons was quantified using a bespoke script developed in Fiji (version 2.14). ^16,19^

### Western blot

Neurons were washed with ice-cold PBS before harvesting in RIPA buffer with phosphatase and protease inhibitors. In cases of RA serum/SF stimulation, neurons were washed twice to minimise residual protein carry-over. BCA assays were performed to measure protein concentration to normalise protein for loading.

Samples were mixed with 4x Laemmli buffer and boiled at 95°C for 5 minutes. Samples and a protein ladder were loaded onto Novex™ Tris-Glycine Mini Protein Gels and run at 100V for 1.5 hours. Protein was transferred from the gel to methanol-activated 0.45 µm PVDF membranes using the Trans-Blot Turbo Transfer kit. Membrane blocking was performed at room temperature for 1 hour in 5% skimmed milk, dissolved in Tris-buffered saline with 0.1% Tween® 20 detergent (TBST), and then probed with primary anti-pSTAT3 antibody over night at 4°C. After three 5-minute washes in TBST, membranes were incubated with secondary Ab (diluted in TBST) for 1 hour. After washing, blots were imaged using SuperSignal Chemiluminescent Substrates. Data were acquired by Chemidoc XRS+ and Image Lab software (version 6.1). Blots were stripped in Millipore stripping buffer for 15 minutes, then blocked and incubated with primary anti-STAT3 antibody, following the same steps as above. Densitometric analyses were conducted in Fiji (version 2.14) to calculate pSTAT3/STAT3 ratios.

### RNA extraction, cDNA conversion and qPCR

Neurons were lysed in RLT buffer with beta-mercaptoethanol and RNA extracted using a Qiagen RNeasy Kit. RNA concentration was measured using a Qubit High Sensitivity kit. The same amount of RNA was converted to cDNA using SuperScript III and 2 ng/mL cDNA was used for qPCR using the Roche LightCycler® FastStart DNA Master SYBR Green mix. Primers used in the study are listed in **Supplementary Table 2**. ΔCycling Thresholds (CT) were obtained by normalising CT values to the housekeeping genes. The data are presented as either 2^-ΔCT^ or fold change (ΔΔCT), which was calculated by dividing 2^-ΔCT^ in each condition by the average 2^-ΔCT^ of control.

### Flow cytometry

IPSC were detached from the well using TrypLE, washed with PBS, then stained with fixable Viability Dye for 20 minutes at 4°C. Cells were washed with FACS buffer (1% BSA, 0.1% sodium azide in PBS) at 5900*g* for 3.5 minutes and blocked in 10% human Ab serum in FACS buffer for 10 minutes at 4°C. TRA-1-60 mAb was used to stain IPSC for 30 minutes at 4°C. After a final wash, IPSC were fixed with 2% PFA for 15 minutes at 4°C before washed a final time and resuspended in FACS buffer. Unstained cells and live/dead controls were prepared to optimize fluorochrome acquisition. Unstained cells were taken through the same workflow as stained cells, without the inclusion of antibodies. For live/dead controls, half of the cells were heat-inactivated at 65°C for 5 minutes followed by a cold shock on ice for 5 minutes, then combined with another half of live IPSC and stained with fixable Viability Dye. Samples were acquired using a FACS Canto-II. Data were analysed using FlowJo software (version 10.8).

### Luminex

A Luminex Discovery Assay (R&D) was used to determine the concentration of IL-6, IL-6R alpha, Oncostatin M (OSM), IFN-alpha, IFN-beta, IL-11 and LIF. Samples were diluted 1:2 (cell culture media, serum and SF) and 1:100 (serum and SF). The plate signal was acquired using FLEXMAP 3D and xPonent software. Experiments and analyses were carried out following manufacturer’s instructions. A five-parameter logistic (5-PL) curve-fit was performed to establish standard curves. When the value could not be determined by the standard curve, the highest or lowest value obtained for the analyte was assigned to the sample.

### Multi-electrode array (MEA) recording and analyses

Neurons were replated at day 11 on Axion Cytoview 24-well plates coated with 0.1% PEI (diluted in 1x borate buffer) and 8uL 1/50 Geltrex (diluted in Knockout DMEM), at 50k cells per well. The spontaneous activity of mature neurons (>day 50) was recorded using the Axion Maestro Edge machine (Axion Biosystems) and Axis Navigator software (version 3.9.1). Before the experiment, neurons were acclimatised with starvation media (Neurobasal+glutamax) for at least 4 hours then recorded at baseline. Cytokines or vehicle (0.1% BSA) were added 1 hour later and neurons were recorded every 6 hours for 2 minutes until 90 hours. At 92 hours, a temperature ramp was applied from 37°C to 41.5°C, with 1.5°C increments and held at each step for 2 minutes. Continuous data were acquired simultaneously at 12.5 kHz per electrode and filtered using a 1-pole Butterworth band-pass filter (200–3,000 Hz). Individual spikes were counted if they crossed a 5.5 standard deviation threshold. A well was considered active if there were at least 4 spikes detected in the four temperature steps analysed.

### Statistical analyses

Statistical tests and sensitivity analyses were performed using Microscoft Excel, R (version 4.1.2) or G*Power (version 3.1.9). The statistical tests used were specified in figure legends. Sensitivity analyses can be found in the **Supplementary Methods**. Plots were generated in GraphPad Prism 10 or using ggplot2 in R studio ^20^. Details of all the statistical tests performed in this study (including n numbers, units of analyses and observed effects) are summarised in **Supplementary Table 3**.

## Results

We first interrogated publicly available databases for the expression levels of the components of JAK/STAT signalling in human sensory neurons. Analyses of two human post-mortem dorsal root ganglia (DRG) neuron single-nuclei RNA sequencing studies and one compilation study showed that JAK1 and STAT3 are widely expressed in human sensory neurons, at levels higher than other JAKs and STATs ^2,3,21^ **(Fig. 1A-B, Supplementary Fig. 1A).** We investigated which STAT3-signalling cytokines may be able to induce neuronal pSTAT3 directly, by identifying the transcript levels of their corresponding receptors on post-mortem human sensory neurons **(Fig. 1C).** We found that *IL6ST*, *LIFR*, *OSMR*, *IL31RA*, *IFNAR1* and *IFNAR2* were moderately to highly expressed, highlighting a possible role for IL-6 family cytokines and type-I interferons in sensory neuron activation.

**Fig. 1:**
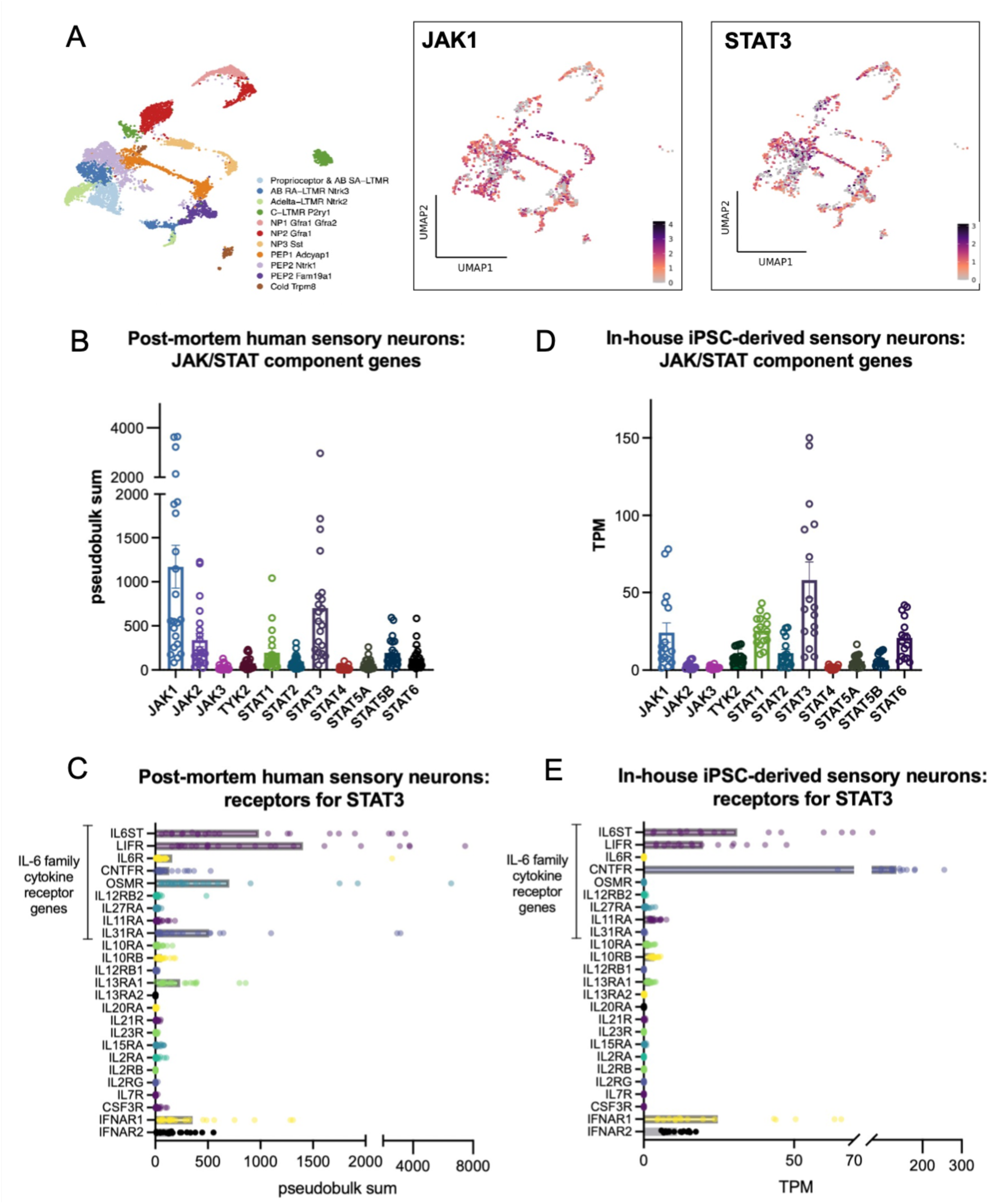
Genes related to JAK/STAT signalling are expressed in human sensory neurons and IPSC-derived sensory neurons. **A.** Single-nuclei RNA sequencing datasets (XSpecies DRG Atlas: http://research-pub.gene.com/XSpeciesDRGAtlas/) ^2^ were analysed for the expression of JAK1 and STAT3 across different subtypes of human post-mortem sensory neurons. Colour scales indicate scaled average expression across samples. **B-E:** Pseudobulk analyses of scRNA-seq of human post-mortem ^2,3^ **(B)** and in-house IPSC-derived sensory neurons for the indicated JAK and STAT molecules **(D).** Expression patterns of the receptors for the indicated STAT3 signalling cytokines in human post-mortem **(C)** and in-house IPSC- derived sensory neurons **(E)**. TPM: transcripts per million.

Since we used human IPSC-derived sensory neurons as a model system in our study, we confirmed an overall similar gene expression pattern of JAK/STAT genes in the neurons we differentiate in-house **(Fig. 1D)**. These results were in keeping with a publicly available atlas of sensory neurons differentiated with a similar protocol ^22^ **(Supplementary Fig. 1B).** Moreover, with some exceptions (*IL31RA*, *OSMR* and *CNTFR),* the expression patterns of JAK/STAT3-relevant cytokine receptors were largely similar to human post-mortem sensory neurons. In particular, high abundance of *IL6ST*, *LIFR*, *IFNAR1* and *IFNAR2* was recapitulated **(Fig. 1E, Supplementary Fig. 1B).**

In order to investigate this further at a cellular and functional level, we derived sensory neurons from two IPSC lines, Kute4 and UKB. We first confirmed that both IPSC lines were in an undifferentiated state using flow cytometry for the marker TRA-1-60 **(Fig. 2A)**. Upon differentiation using the Chambers protocol ^17^, sensory neurons expressed neuronal markers somatostatin (*SST*) and Nav1.7 (*SCN9A*), with glial and fibroblast markers *FABP7* and *COL15A1* largely absent **(Fig. 2B).** At protein level, most of the neurons were positive for BRN3A, a sensory neuron transcription factor, and PGP9.5, a pan-neuronal marker, with a purity of over 90% across all the batches used in this study **(Fig. 2C-D).** These data indicate that our IPSC-derived sensory neurons were pure, with a phenotype similar to human sensory neurons, therefore suitable for studying STAT3 signalling.

**Fig. 2:**
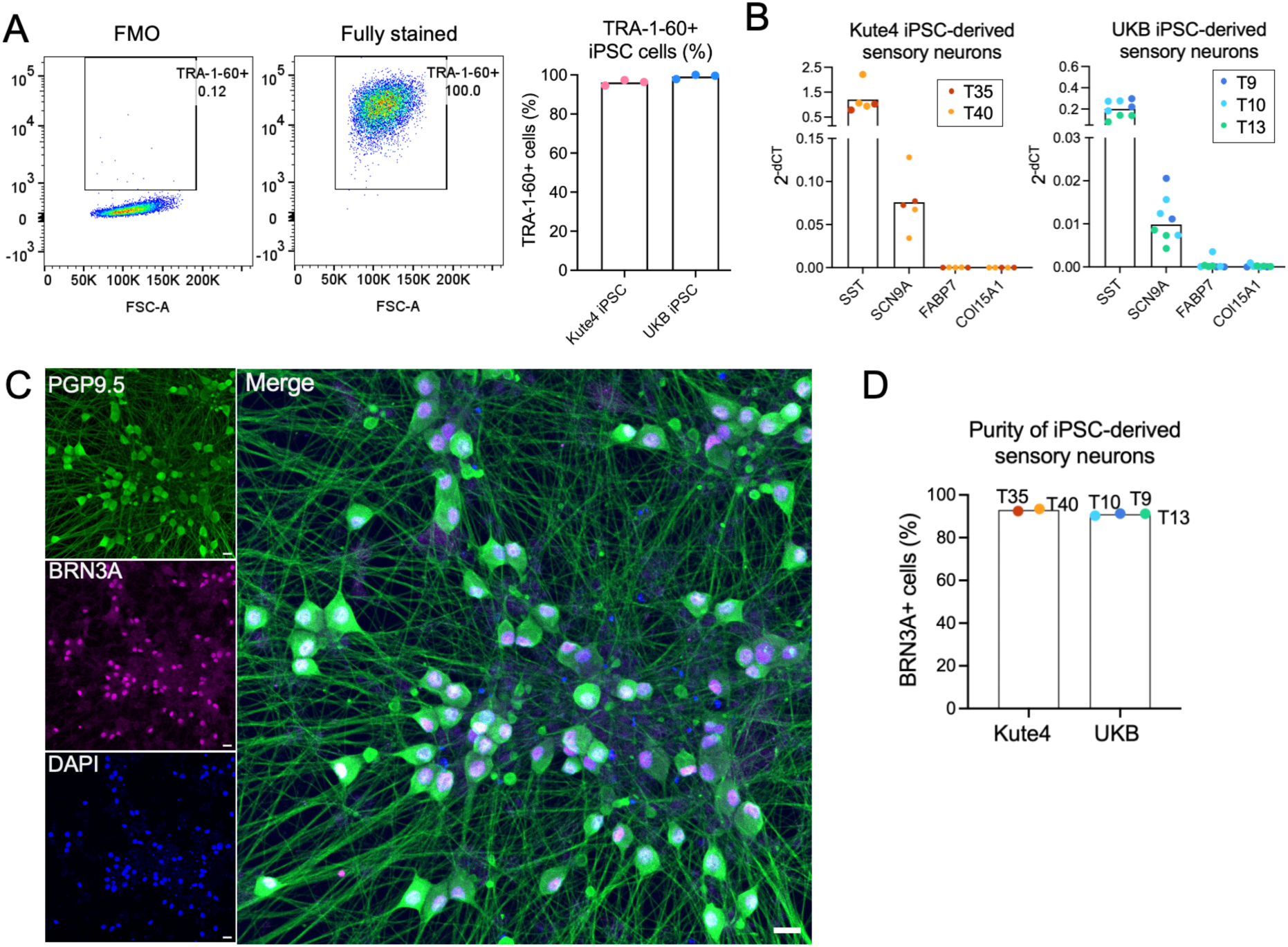
Characterisation of IPSC-derived sensory neurons. **A.** IPSC-derived sensory neurons were differentiated from Kute4 and UKB lines and stained for the expression of TRA-1-60, a marker of undifferentiated cells. Representative flow cytometry plots and cumulative data (n=3) showing the percentage of TRA-1-60+ cells in UKB IPSC at passage 14, as compared to the fluorescence minus one (FMO) control (left panel). **B.** Bar plots showing gene expression of neuronal markers (*SST*, *SCN9A*) and non-neuronal genes (*FABP7*, *COL15A1*) by sensory neurons derived from Kute4 (left) and UKB (right) IPSC lines. Within each independent differentiation batch (indicated by trial (T) number and different colours) n=2-3 biological repeats were used. **C, D.** IPSC-derived sensory neurons were differentiated to over day 50 and stained for the sensory neuron marker BRN3A and pan-neuronal marker PGP9.5. Representative immunohistochemistry of day 81 neuron staining shown in (**C**, scale bar: 20 μm) and cumulative data showing the purity of the neuronal culture, quantified as BRN3A+ cells within DAPI+ cells (**D**, n=5 independent differentiation batches of Kute4 and UKB IPSC lines).

We tested whether neuronal STAT3 expression was functionally relevant by investigating its ability to become phosphorylated (pSTAT3, Tyr705) in the context of the inflamed RA joint environment. For this, we incubated IPSC-derived sensory neurons with a panel of paired serum and joint-derived synovial fluid (SF) donated by individuals with RA. We found that several RA SF samples were able to induce neuronal pSTAT3 albeit to varying degrees, while paired serum did not **(Fig. 3A, Supplementary Fig. 2A).** This suggests that neuronal pSTAT3 is not induced by a common feature of human biofluid but is more likely driven by specific inflammatory cytokines enriched in RA SF. Moreover, RA SF-induced pSTAT3 co-localised with BRN3A, confirming signalling specific to sensory neurons **(Supplementary Fig. 2B).** To ascertain the role of the JAK/STAT pathway in these findings, we repeated our experiments in the absence or presence of tofacitinib, a clinically approved JAKi. We observed a complete block of neuronal pSTAT3 induction by RA SF in the presence of tofacitinib **(Fig. 3B-C).** Taken together, our data support the possibility that JAK/STAT3 signalling is activated in sensory neurons in some individuals with RA and that JAKi can directly inhibit that activation.

**Fig. 3.**
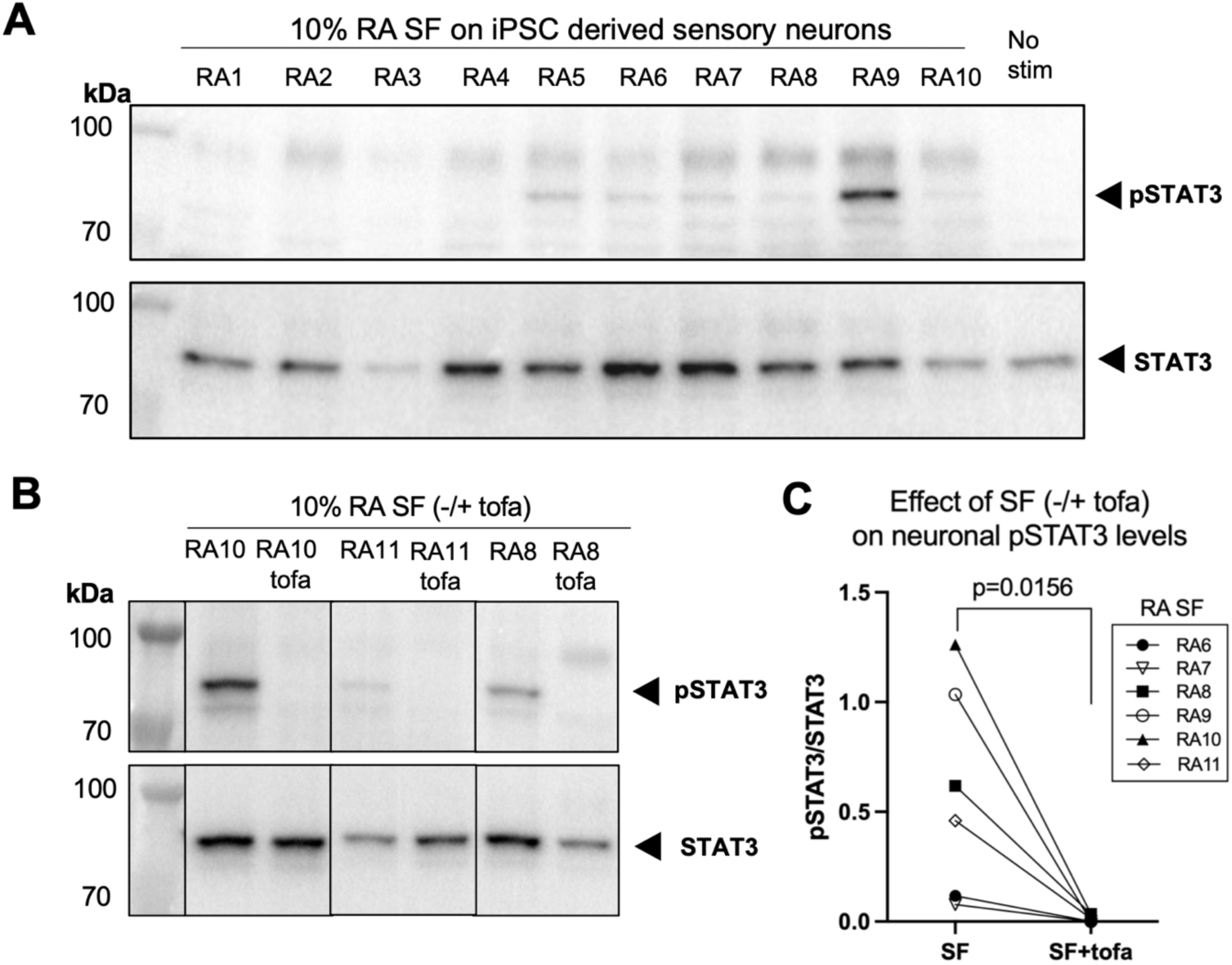
RA-derived synovial fluid can induce pSTAT3 in human sensory neurons cultures, which is blocked by tofacitinib. **A.** IPSC-derived sensory neurons (day 55) were incubated for 1 hour with 10% RA SF (n=10) or media alone (No stim). Cells were lysed and probed for pSTAT3 (upper panel) and STAT3 (lower panel) using Western blot. **B, C.** IPSC-derived sensory neurons were stimulated as in (**A**) in the absence or presence of tofacitinib (2μM) for 1 hour and pSTAT3/STAT3 expression analysed. Representative Western blot (**B**) and cumulative data (**C**) showing the quantification of the ratio of pSTAT3/STAT3 signal of neurons from two batches (day 55-65) stimulated with 10% RA SF from n=6 different individuals with RA in the absence or presence of tofacitinib. Data were analysed using a paired nonparametric t-test.

We next sought to determine which cytokines may contribute to the observed neuronal pSTAT3 induction by RA SF. Given that induction was specific to SF (vs. patient-matched serum), we were particularly interested in identifying JAK/STAT3-relevant cytokines that are enriched in SF. We focused our experiment on cytokines with corresponding receptors on sensory neurons, specifically type-I interferons (IFN-alpha, IFN-beta) and IL-6 family cytokines: LIF; IL-6 and IL-11, which can signal through IL-6ST (gp130) with their corresponding soluble receptors; and OSM, which can signal through the LIF receptor. We compared the levels of these cytokines in 12 paired serum and SF samples from patients with seropositive RA and 4 healthy control serum samples using Luminex **(Supplementary Table 4).**

We found elevated levels of IL-6, IL-11, LIF, IFN-alpha and IFN-beta in RA SF compared to RA serum, whilst OSM levels were high in both. Among these cytokines, IL-6 and IL-11 were particularly enriched in SF **(Fig. 4).** While sensory neurons do not express membrane-bound IL-6 receptor **(Fig. 1C&E)**, soluble IL-6 receptor (IL-6Ra) was abundantly detected in all the serum and SF samples we investigated, in agreement with a previous report ^23^ **(Fig. 4, Supplementary Table 5)**. This suggests that trans-signalling of IL-6 with its co-receptor gp130 could take place on neurons. We also noted that the levels of IL-6, LIF and IFN-alpha in RA SF correlated well with RA SF-induced neuronal pSTAT3 levels as measured by Western blot **(Supplementary Fig. 3)**.

**Fig. 4.**
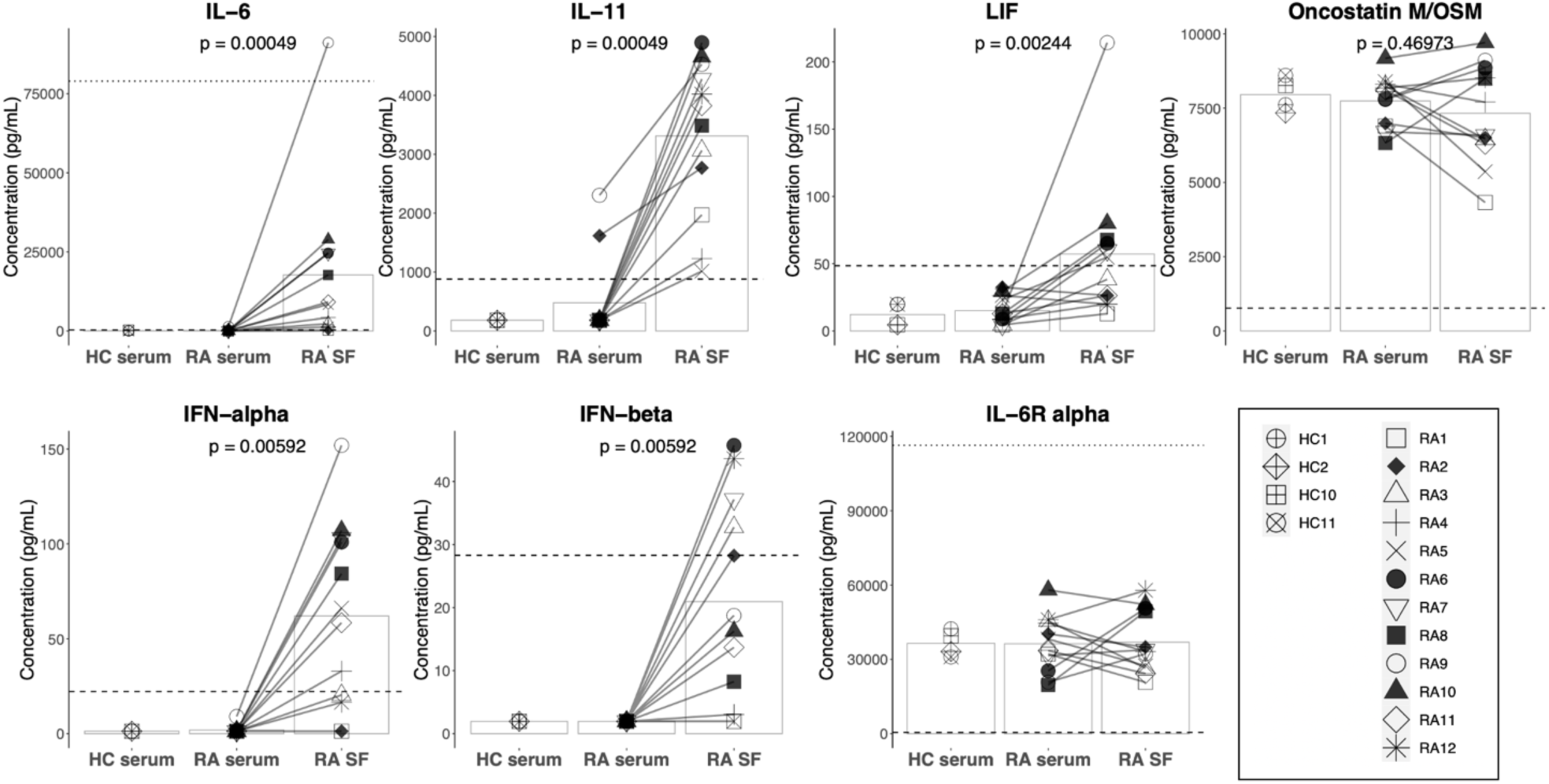
Cytokines upstream of STAT3 signaling are enriched in RA SF. Healthy control (HC) serum (n=4) and paired RA serum and SF (n=12) were analysed for the presence of IL-6, IL-11, LIF, OSM, IFN-alpha, IFN-beta and IL-6R alpha by Luminex assay. Statistical significance was assessed using paired nonparametric t-tests (RA SF vs. serum); p < 0.007 was considered significant after Bonferroni correction for the six cytokines and indicated by *. The dotted lines for IL-6 and IL-6R alpha graphs indicate the highest standard value. The dashed lines indicate the lowest standard value. The legend symbols are representative of individual HC or RA samples; the bars represent the data means. Standard values, sensitivity, and all individual means (+SD) of the assay are listed in **Supplementary Table 5**.

To identify which cell types in RA synovial tissue might express these cytokines, we examined sequencing data from the Accelerating Medicines Partnership (AMP) RA Phase-1 initiative ^24^. Fibroblasts, T cells, B cells and monocytes all expressed mRNA for IL-6, LIF, OSM, as well as IL-6 receptors and IL-11 receptors **(Supplementary Fig. 4).** To confirm these findings at protein level, we set up independent cultures of RA synovial fibroblasts or RA-derived mononuclear cells and tested their cell culture supernatant for the presence of these cytokines by Luminex. When RA synovial fibroblasts were stimulated with IL-1beta ^25^, they became a rich source of IL-6, IL-11, LIF and IFN-alpha **(Supplementary Fig. 5A)**. Similarly, when peripheral blood mononuclear cells (PBMC) and synovial fluid mononuclear cells (SFMC) were subjected to T cell stimulatory conditions (anti-CD3 + anti-CD28 mAbs), they increased the expression of IL-6, LIF and IFN-alpha **(Supplementary Fig. 5B)**. These data support the notion that activated fibroblasts and immune cells present in inflamed RA joints can be a cellular source of STAT3 signalling cytokines capable of activating sensory neurons directly.

We next asked whether SF induction of neuronal pSTAT3 could be recapitulated with recombinant cytokines, namely IL-6 +/- soluble IL-6R (sIL-6R), LIF, IL-11 +/- soluble IL-11R (sIL-11R), IFN-alpha and IFN-beta. All five mediators induced neuronal pSTAT3 at the time points investigated **(Fig. 5 A-D)**. Using IL-6+sIL-6R and LIF as examples, we assessed whether the pSTAT3 signal could be blocked by tofacitinib. We found that tofacitinib and two other clinically approved JAKi, baricitinib and updacitinib, completely blocked IL-6+sIL-6R or LIF-induced neuronal pSTAT3 **(Fig. 5E, Supplementary Fig. 6).** To ensure that the pSTAT3 signal observed by Western blot was derived from neurons and not the occasional non-neuronal cell, we used immunocytochemistry to confirm co-localisation of pSTAT3 with neuronal markers BRN3A and NF200 following stimulation with IL-6+sIL-6R or LIF **(Fig. 5F).** Taken together, these data indicate that cytokines enriched in the inflamed RA joint can directly induce neuronal pSTAT3, and that JAKi can block such effects.

**Fig. 5.**
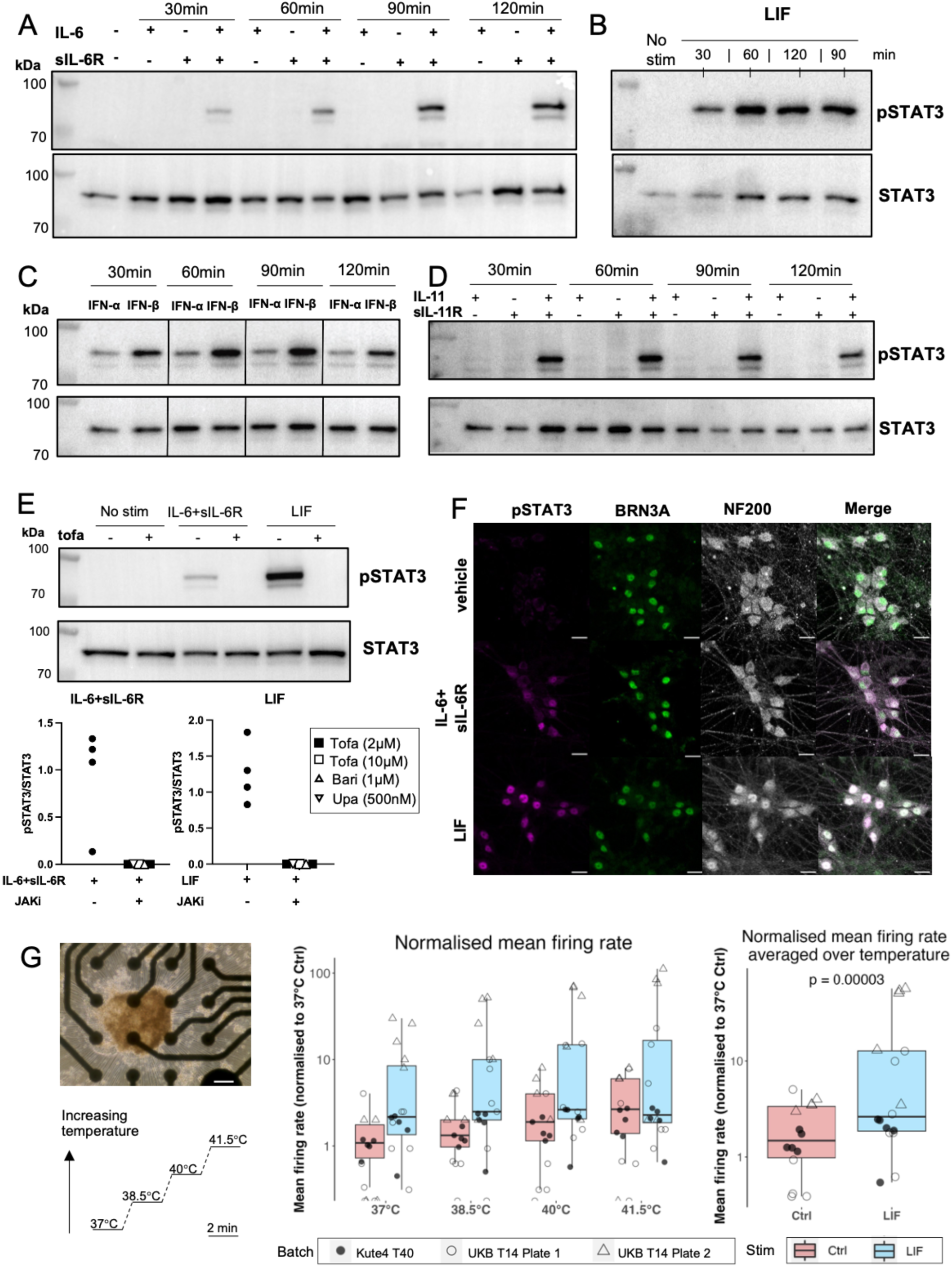
RA-relevant recombinant cytokines induced neuronal pSTAT3 and elevated firing rates in peripheral sensory neurons. **A-E**: IPSC-derived sensory neurons were incubated for 30, 60, 90 and 120 minutes with human recombinant **(A)** IL-6, sIL-6R or both (100ng/mL), **(B)** LIF (100ng/mL), **(C)** IFN-α (300U/mL), IFN-β (300U/mL), or **(D)** IL-11, sIL-11R or both (100ng/mL). Cell lysates were probed for pSTAT3 and STAT3 by Western blot. **E.** Neurons from two batches were incubated with IL-6+sIL-6R or LIF without or with JAKi for 1 hour. Representative blot of neurons blocked using 10μΜ tofacitinib and cumulative data using tofacitinib, baricitinib or updacitinib (n=4-5). **F.** Neurons (day 77) were stained for pSTAT3 and sensory neurons markers BRN3A and NF200 by immunocytochemistry, shown in individual or merged channels, scale bar: 20μm. **G.** Neurons plated on an MEA plate (scale bar: 20μm) treated without or with 100ng/mL LIF for 92 hours, and subjected to increased temperature. Boxplots of Ctrl (red) and LIF (blue) showing normalised mean firing rates at each temperature (middle) or averaged across temperature (right). The p-value derives from a nonparametric t-test to assess the effect of LIF. Legend symbols represent wells from three plates and two IPSC lines (n=14-16, neuronal age: day 64-75).

Since LIF is a lesser-known pain mediator compared to IL-6, as a tentative test we assessed whether LIF could affect neuronal excitability. Acute application did not increase neuronal firing rate initially; however, an increase was observed over time, at around 92 hours (**Supplementary Fig. 7A-B**). At this timepoint, we subjected the neurons to a temperature gradient ramping up from 37°C to 41.5°C (**Fig. 5G, Supplementary Fig. 7C-D**). As expected, neurons showed an overall trend in increasing their firing rate as temperatures increased, although this was not statistically significant. However, we observed a significant main effect of LIF (repeated measures ANOVA, p=0.045) in modulating neuronal firing rate across the temperature range. This was even more apparent when mean firing rates were averaged over the temperature steps (p=0.00003).

As a final step, we explored the downstream intracellular consequences of STAT3 activation in neurons. We re-analysed a publicly available chromatin-immunoprecipitation sequencing (ChIP-seq) dataset generated in mouse DRG neurons to identify possible pSTAT3 binding sites of four genes of interest ^1^ **(Fig. 6A)**. As expected, sensory neurons showed pSTAT3 binding in the promoter region of SOCS3, the negative regulator of STAT3 activation ^26^. Of interest to sensory neuron function, there also appeared to be pSTAT3 binding sites in several well-known pain-relevant genes: ATF3, an injury and regeneration marker ^27,28^; BDNF, a growth factor that mediates central sensitization ^29,30^; and CSF1, a cytokine that promotes microglia and macrophage activation in DRG and spinal cord, likely exacerbating pain ^31,32^. To confirm involvement of these genes in our human IPSC-derived sensory neuron system, we probed for changes in their gene expression after 24h stimulation with IL-6+sIL-6R **(Fig. 6B-C).** An increase in expression was observed for all four genes, as would be expected upon pSTAT3 binding. A similar and even more robust pattern was seen with LIF stimulation and importantly, transcriptional regulation was entirely reversed by the addition of a JAK inhibitor **(Fig. 6C)**. Together, these data reveal that JAKi can directly dampen upregulation of pain-relevant genes in sensory neurons, pointing towards a potential mechanistic explanation for why JAKi are particularly effective at reducing pain in RA.

**Fig. 6.**
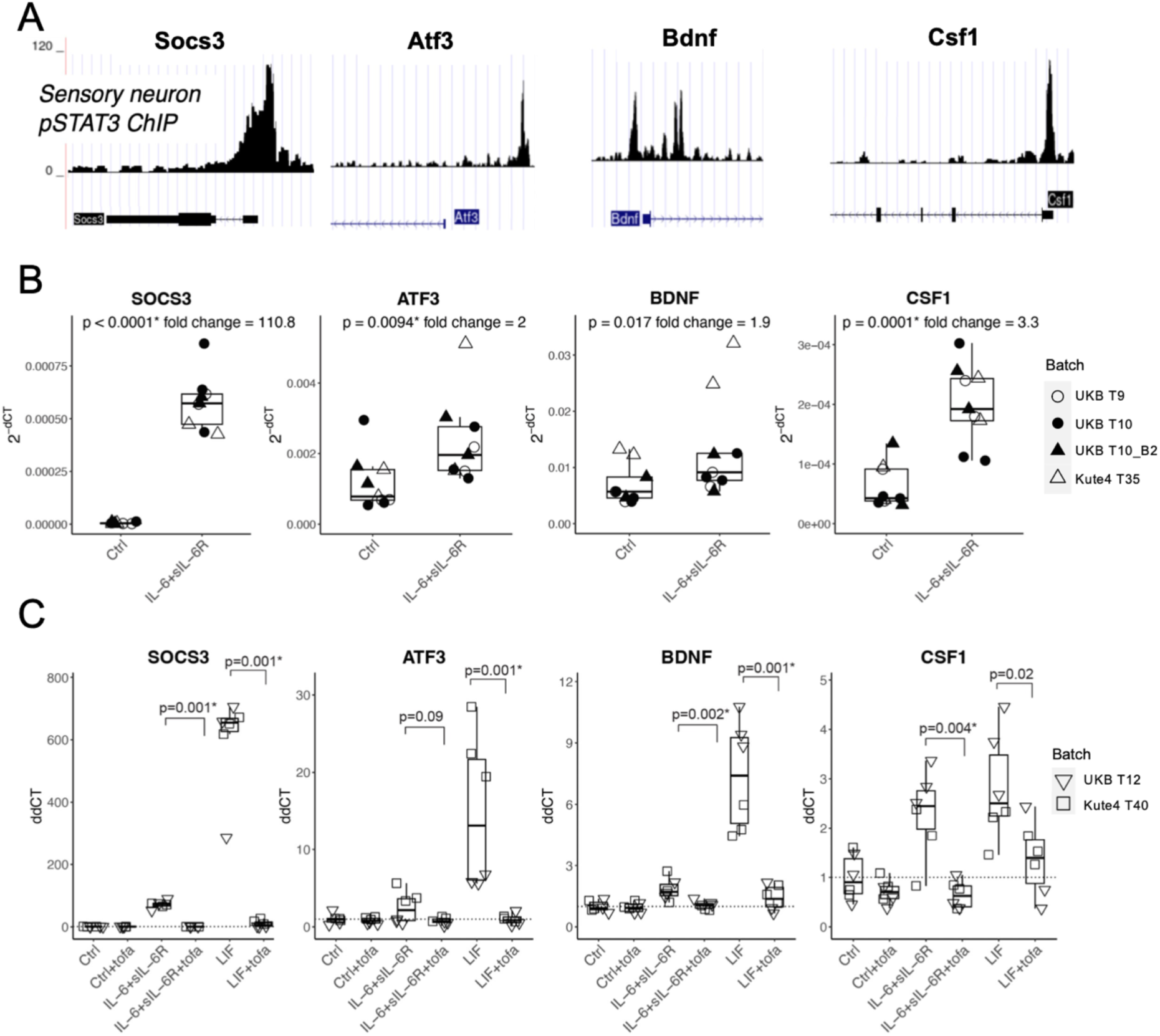
IL-6+sIL-6R and LIF increase the expression of pain-relevant genes with pSTAT3 binding sites, which is reversed by JAKi. **A.** ChIP sequencing dataset ^1^ showing pSTAT3 binding sites in *SOCS3*, *ATF3*, *BDNF* and *CSF1* in mouse DRG neurons. **B.** IPSC-derived sensory neurons were treated without or with 100 ng/mL IL-6+sIL-6R for 24 hours, followed by qPCR analysis. Boxplots showing gene expression of the indicated genes. Data derived from n=9 culture wells using three independent differentiations (designated by the prefix “T”) from two IPSC lines (day 67-76). **C.** IPSC-derived sensory neurons were treated without or with IL-6+sIL-6R or LIF (all at 100 ng/mL) for 24 hours in the absence or presence of tofacitinib (2μM), followed by qPCR analysis. Box plots showing expression of the indicated genes. Data are derived from n=6 neuronal wells derived from two independent IPSC lines (day 57-61). Statistical significance was determined by one-tailed nonparametric t-tests to compare between with or without IL-6+sIL-6R (**B**) or with or without tofacitinib in the presence of IL-6+sIL-6R or LIF (**C**). Neurons aged day 57-76 were used. p < 0.013 was deemed significant after Bonferroni correction and marked by *.

## Discussion

JAKi are frequently used in the treatment for RA due to their effectiveness, affordability, and convenience of administration ^8^. In light of evidence that JAKi may be more effective than anti-TNF in reducing pain in RA ^10,11^, we tested the hypothesis that JAKi have a direct inhibitory effect on JAK/STAT signalling in human IPSC-derived sensory neurons.

We showed that RA SF taken from the inflamed joints of individuals with RA can induce neuronal pSTAT3, which can be blocked by JAKi. We present evidence that this effect is likely due to the presence of IL-6 family cytokines (including IL-6, IL-11, LIF) and/or Type-1 interferons in the SF. We further showed that the corresponding recombinant cytokines recapitulated neuronal pSTAT3 induction, which was blocked by JAKi. Some of these cytokines (IL-6, IFN-alpha) are already known pain mediators ^33–36^. We provided initial evidence that LIF is another cytokine that can induce neuronal sensitisation. Finally, the activation of pSTAT3 induced by IL-6+sIL-6R or LIF increased the expression of pain-related genes in IPSC-derived sensory neurons, which was blocked by JAKi. We hence propose that JAKi may have dual action, dampening both inflammation and neuronal activation in RA joints.

Our study adds to the pool of knowledge on the presence and abundance of inflammatory cytokines in RA SF and serum. Consistent with previous reports, we found a higher concentration of IL-6, IL-11, LIF and IFN-alpha in RA SF compared to paired serum ^37–41^. To our knowledge, this is the first time that levels of IFN-beta and OSM were measured and compared in HC serum, RA serum and paired SF. The level of OSM was high and similar across all samples while IFN-beta was elevated in RA SF compared to RA serum.

Our study presents evidence of a direct effect of some of these cytokines (IL-6, LIF, IL-11 and type 1 interferons) on sensory neurons. This finding is corroborated by a recent publication that showed that IL-6+sIL-6R can elevate *SOCS3* and *CSF1* in a mouse neuroblastoma cell line ^42^. In keeping with our results, the authors also reported that *SOCS3*, *ATF3* and *CSF1* were elevated in sensory ganglia in a mouse model of arthritis, an effect that could be reduced by intragastric injection of baricitinib.

Of the JAK/STAT cytokines we studied, IL-6 and type-1 interferon are already known to be able to elevate neuronal excitability *in vivo* and in primary mouse DRG neurons ^33–35^. Here, we provide initial evidence that the same is true for LIF, which appeared to be able to modulate the firing rate of IPSC-derived sensory neurons in response to temperature. We have yet to test whether RA SF can affect firing rates in a similar fashion. Increased excitability has been observed when mouse DRG neurons were incubated with SF taken from individuals with osteoarthritis (OA), which is also enriched in cytokines such as IL-6 and IL-11 ^43,44^. We would therefore postulate that RA SF will exert similar effects on our human neurons.

Our findings are limited by the fact that the RA SF samples we used were all collected from female patients with active, seropositive RA. While this represents a very prevalent population amongst individuals with RA, it does not reflect the full heterogeneity of those affected by the disease. Whether a similar pattern is seen in seronegative RA, in male patients, or in those with low disease activity remains to be tested. Moreover, for future studies, it would be informative to capture information on pain scores and correlate them with JAK/STAT3 signalling in sensory neurons.

Indeed, with pain being one of the top concerns for individuals living with RA, it is important to understand exactly which JAK/STAT cytokines within RA SF are the most potent modulators of sensory neuron activity. Here we identified multiple candidates that could be exploited in drug development, e.g. one could consider using bispecific antibodies capable of trapping multiple IL-6 family cytokines at once ^45^. Alternative approaches to interfering with JAK/STAT signalling in RA are going to be especially important, given the recent safety concerns regarding the use of JAKi ^46^.

Future work in this area will not only be relevant to RA, but also to other immune-mediated inflammatory diseases where the JAK/STAT pathway is known to be involved in pathogenesis, such as psoriatic arthritis and inflammatory bowel disease^47^. Our work raises the possibility that in all these instances, inhibiting JAK/STAT3 signalling might limit sensory neuron hyperexcitability not only indirectly by dampening inflammation, but also directly by interfering with intracellular pathways in neurons. We hope that this mechanistic insight can accelerate the development of novel painkillers to help improve the quality of life of the millions of individuals currently living with pain induced by RA and other, pathologically similar conditions.

## Supporting information

Western blots raw files

## Acknowledgements

We thank Willow Hight-Warburton for her assistance with Western blot experiments. We thank Cynthia Bishop from Advanced Cytometry Platform for her help with running the Luminex assay and Carl Hobbs for the reagents for pSTAT3 staining. We thank Adam Pavlinek and Deepak Srivastava for helping and providing access to the MEA equipment. We thank James Galloway for his suggestions for this project.

The UKB-GCamP6f line was generated as part of an Innovative Medicines Initiative 2 Joint Undertaking under Grant Agreement no.116072. This Joint Undertaking has received the support from the European Union’s Horizon 2020 research and innovation.

We acknowledge King’s College London as the source of HPSI0714i-kute_4 human iPSC line which was generated under the Human Induced Pluripotent Stem Cell Initiative funded by a grant from the Wellcome Trust and Medical Research Council, supported by the Wellcome Trust (WT098051) and the NIHR/Wellcome Trust Clinical Research Facility, and acknowledge Life Science Technologies Corporation as the provider of CytoTune. This research was supported by Research and Development, Guy’s and St Thomas’ NHS Foundation Trust. The views expressed are those of the author(s) and not necessarily those of the NHS.

## Author contributions

YL conducted the experiments and drafted the manuscript. EHG and RR conducted immune cell and fibroblast experiments for culture supernatant generation. LJK, IZ, AL & LF helped with growing IPSC-derived sensory neurons. RJM supported Western blot experiments. SR processed and catalogued the serum, blood and synovial fluid samples. PR and OB generated the UKB-GCamP6f IPSC line. SJ provided intellectual support. FD & LST conceptualised and co-supervised the project, helped draft the manuscript and provided intellectual support. All authors edited and/or reviewed the final draft of the manuscript.

## Supplementary figures

**Supplementary Fig. 1.**
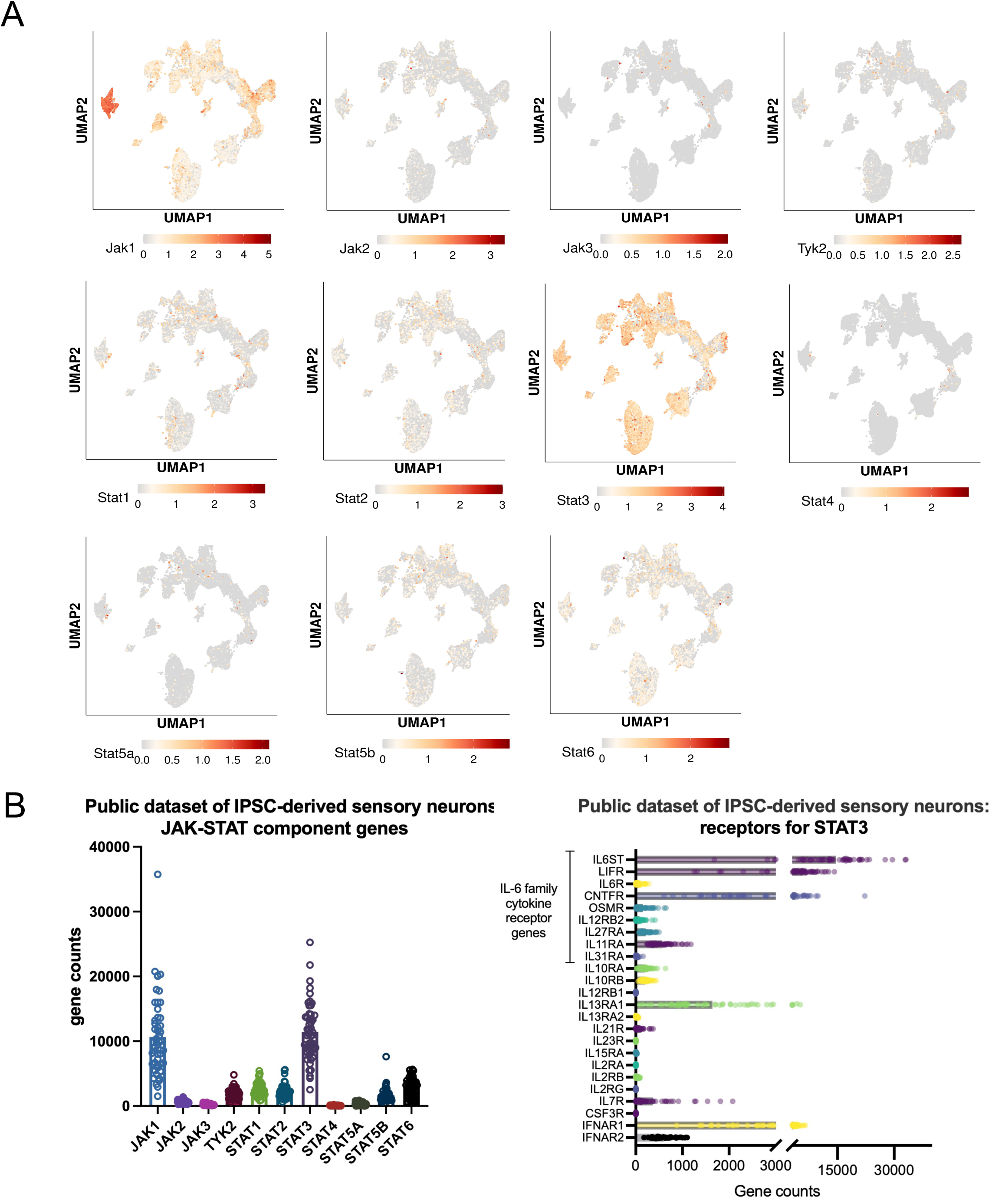
JAK1 and STAT3 are the two most highly expressed components in human and iPSC sensory neurons. **A.** Analyses of a compilation atlas also showed that JAK1 and STAT3 are abundantly expressed by human sensory neurons compared to other JAKs and STATs (18). **B.** A public RNA-seq dataset of iPSC-derived sensory neurons (19) shows very similar expression patterns of JAK/STAT signalling components compared to our in-house iPSC derived sensory neurons.

**Supplementary Fig. 2:**
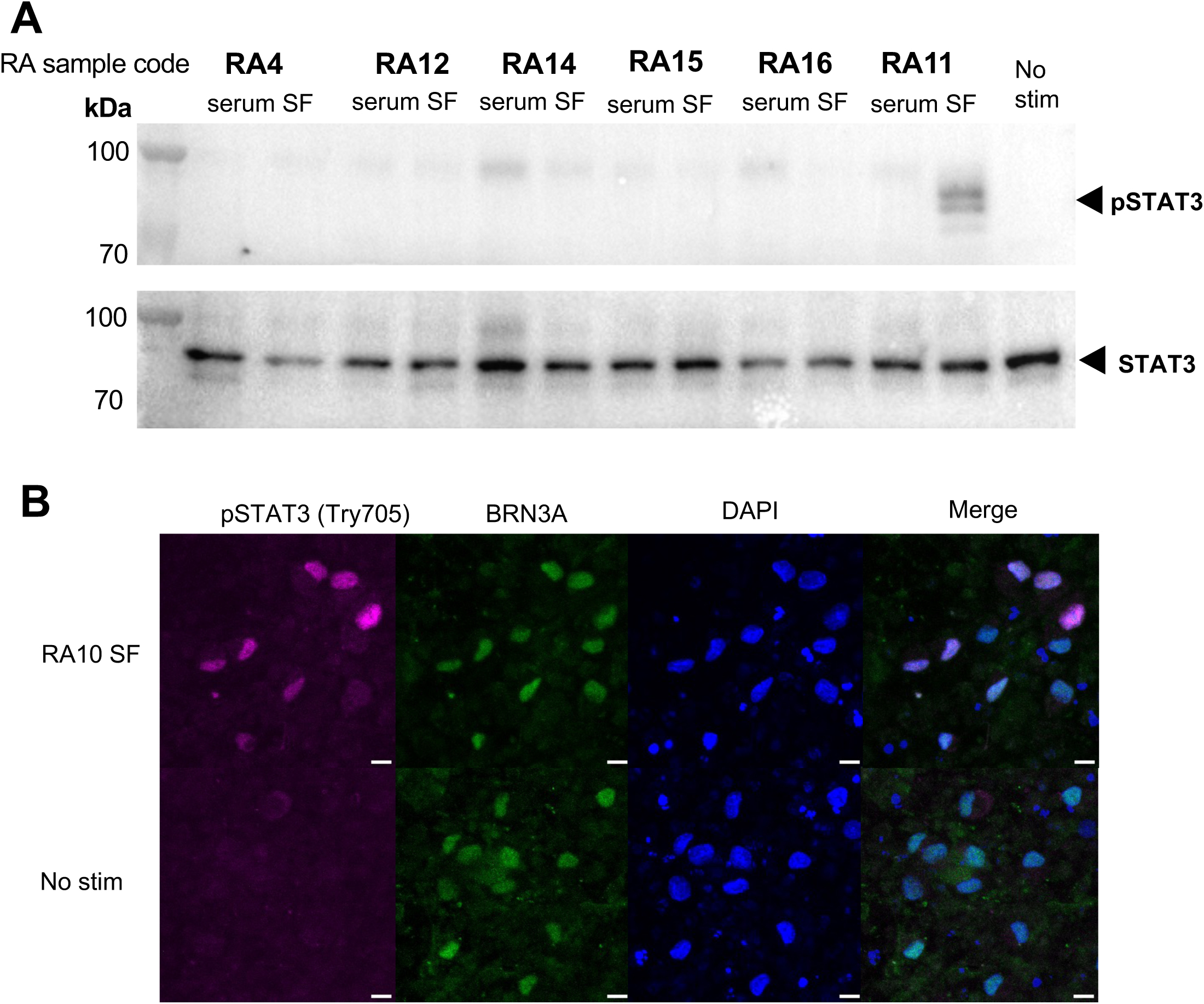
RA SF can induce neuronal pSTAT3. A. RA SF but not paired serum induced pSTAT3 in iPSC-derived sensory neurons cultures. Both SF and serum were used at 10% and applied to sensory neurons for 1 hour before neurons were lysed for Western blot. B. Immunocytochemistry of human iPSC-derived sensory neurons stimulated with 10% SF or media only for 1 hour. There is a strong pSTAT3 signal in the nuclei of more than half of the sensory neurons following 1 hour stimulation. Scale bar: 10µm.

**Supplementary Fig. 3.**
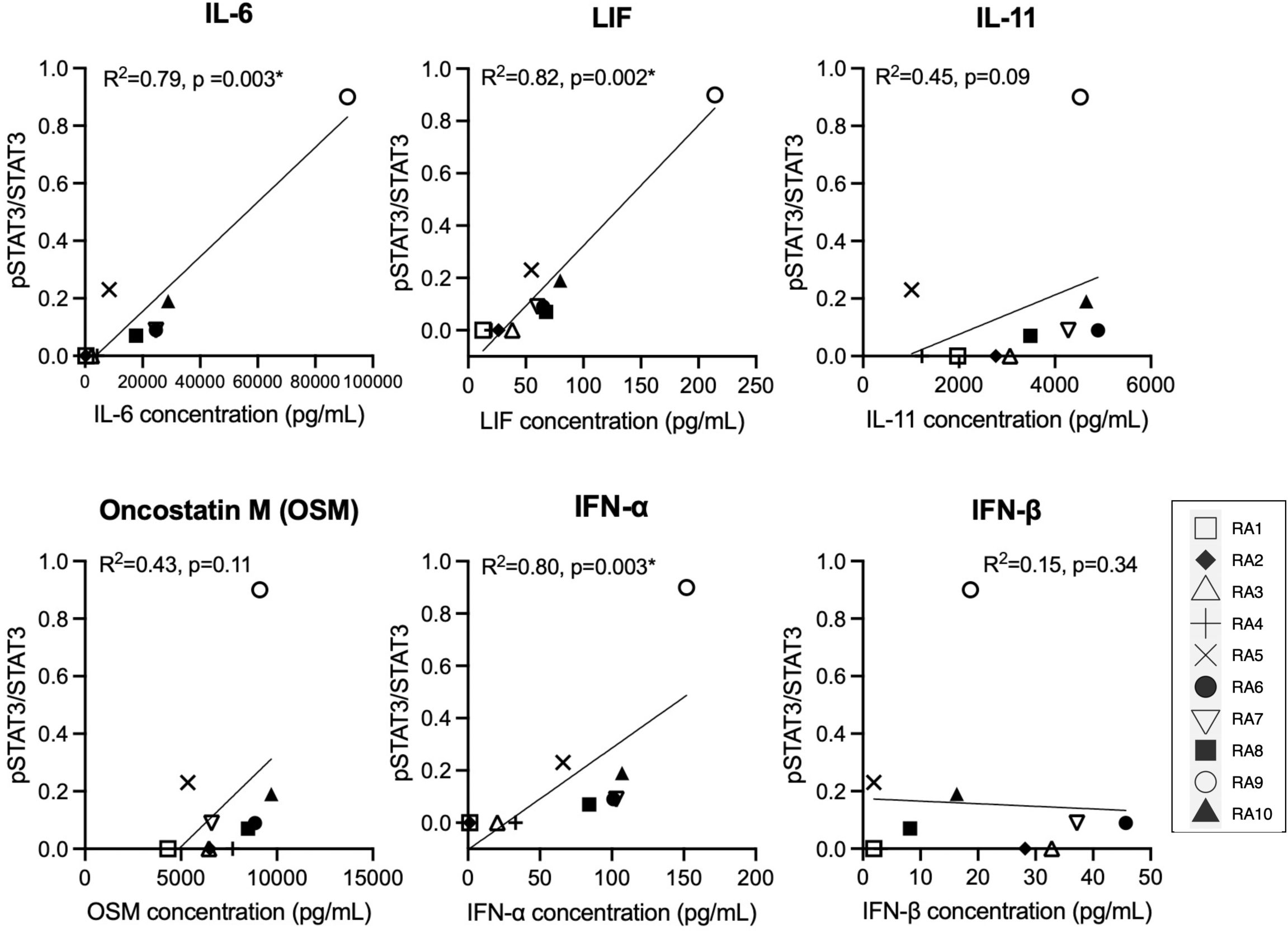
Neuronal pSTAT3 levels induced by RA SF correlated with the cytokine levels in RA SF. Spearman’s correlations of the level of STAT3 cytokines in RA SF (shown in Fig.4) and the neuronal pSTAT3/STAT3 they induced (n=10). Significance after multiple comparison correction indicated by * (cut-off: p=0.008 for 6 cytokines).

**Supplementary Fig. 4.**
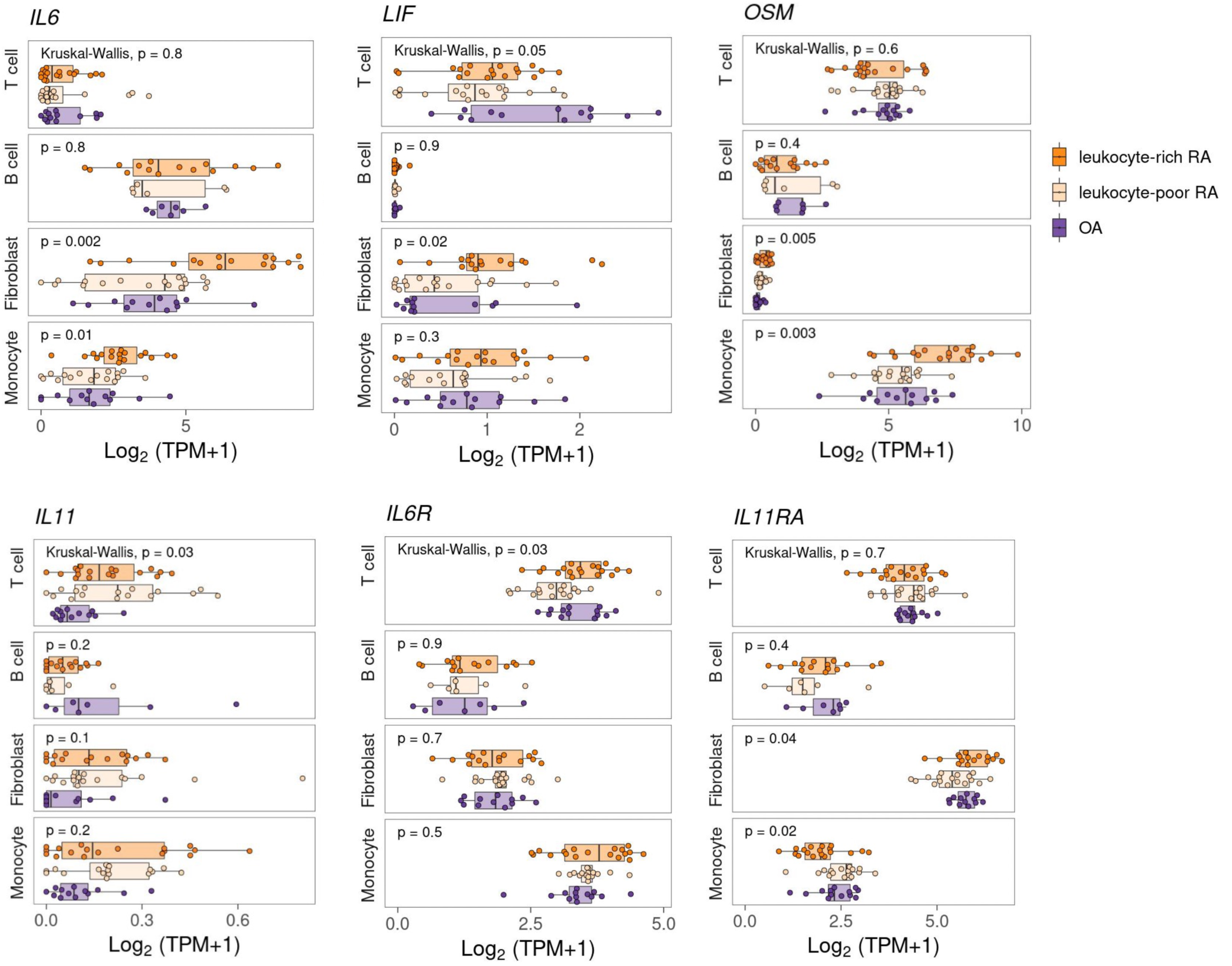
Putative cellular source of STAT3 cytokines. AMP-1 bulk RNA-seq analyses reveal immune cells and fibroblasts produce IL-6, LIF, IL-11, OSM, IL-6R and IL-11RA at mRNA level (21) (https://immunogenomics.io/ampra/).

**Supplementary Fig. 5:**
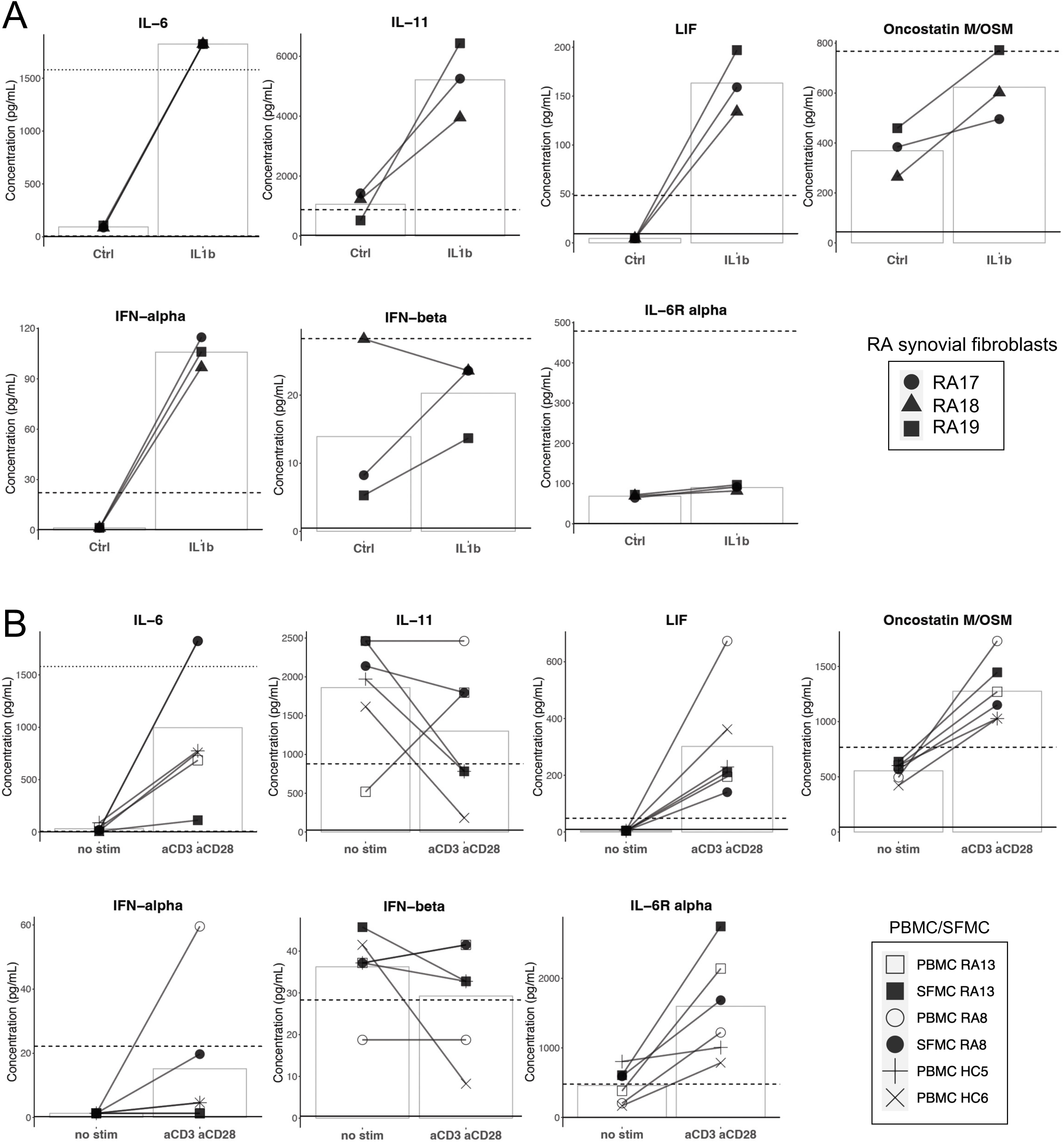
RA synovial fibroblasts and RA PBMC and SFMC are putative sources of IL-6, IL-11, LIF and IFN-alpha. **A.** RA synovial fibroblasts showed increased expression of IL-6, IL-11, OSM and IFN-alpha following 10ng/mL IL-1β stimulation for 24 hours. N = 3 samples from three individuals with RA. Passage 5-6 fibroblasts were used in the experiments. **B.** PBMC and SFMC showed increased expression of IL-6, LIF, OSM following T cell stimulation (1.25 µg/mL anti-CD3 & 1 µg/mL anti-CD28) for 3 days. N = 2 paired samples from two individuals with RA, n=2 samples from HC donors.

**Supplementary Fig. 6.**
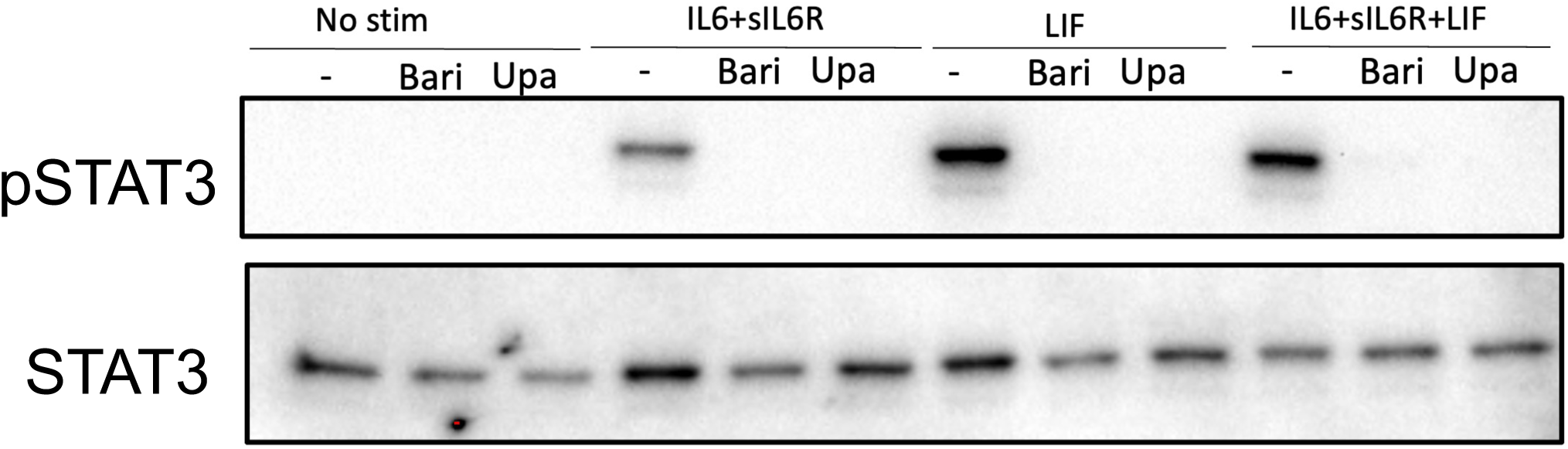
Neuronal pSTAT3 can be blocked by other clinically approved JAKi. Pre-incubation of neurons with baricitinib (1uM) and upadacitinib (0.5uM) for 1 hour completely blocked neuronal pSTAT3 induced by IL-6+sIL-6R and/or LIF (IL-6, sIL-6R, LIF: 100 ng/mL).

**Supplementary Fig. 7:**
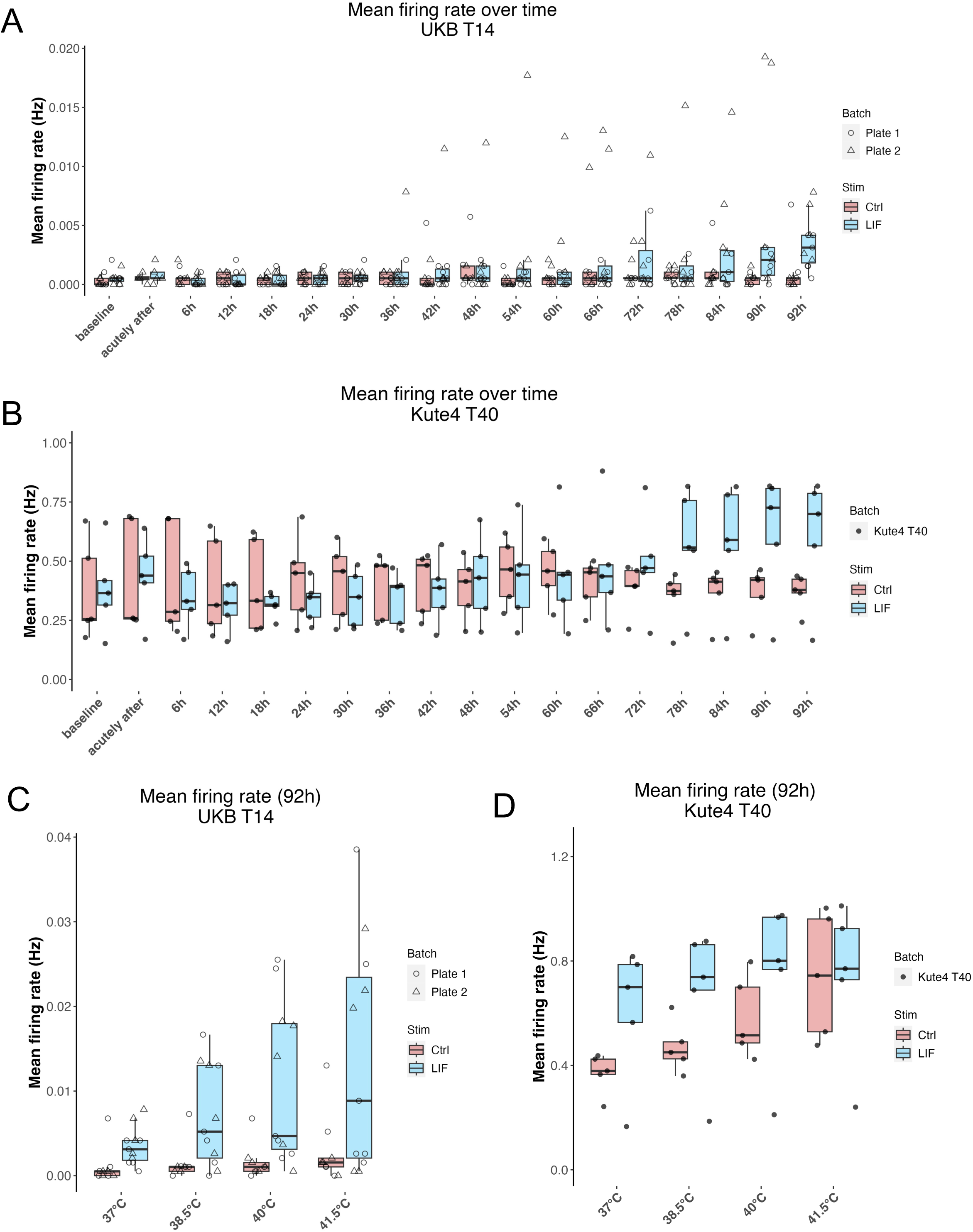
Multi-electrode array analyses of iPSC-derived sensory neurons over time and across temperature steps. **A-B:** Longitudinal mean firing rate of UKB T14 neurons (**A**) and Kute4 T40 neurons (**B**) recorded over 92 hours. **C-D:** Boxplots showing mean firing rate of UKB T14 neurons (**C**) and Kute4 T40 neurons (**D**) at temperature steps from 37°C to 41.5°C. Neuronal age was D60, D67 (UKB T14 Plate 1&2) and D71 (Kute4 T40) at baseline. UKB T14 Plate 1&2: Ctrl (n=9) vs LIF (n=11). Kute4 T40 Ctrl (n=5) vs LIF (n=5).

## Supplementary tables

**Supplementary Table 1.**
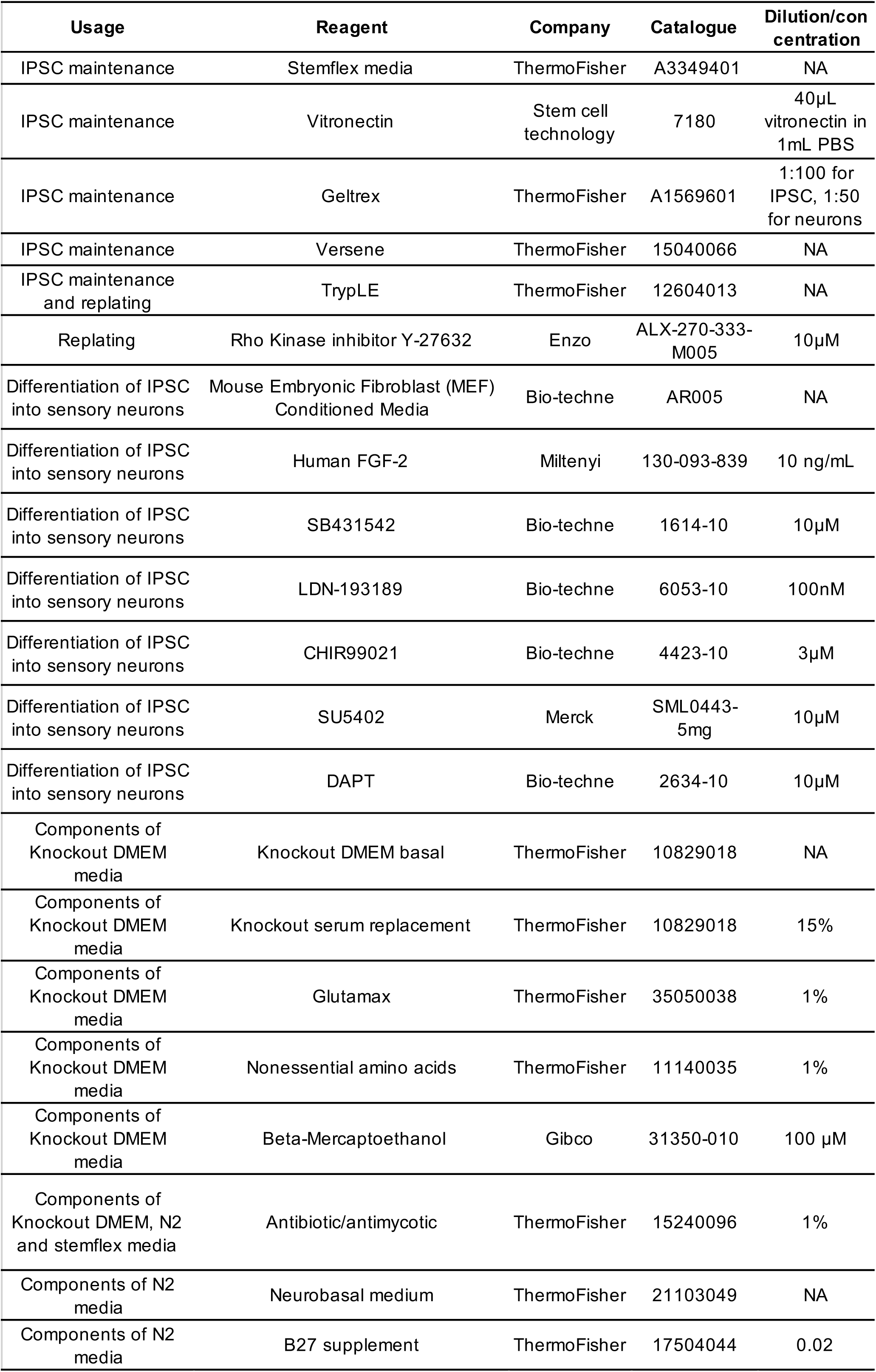
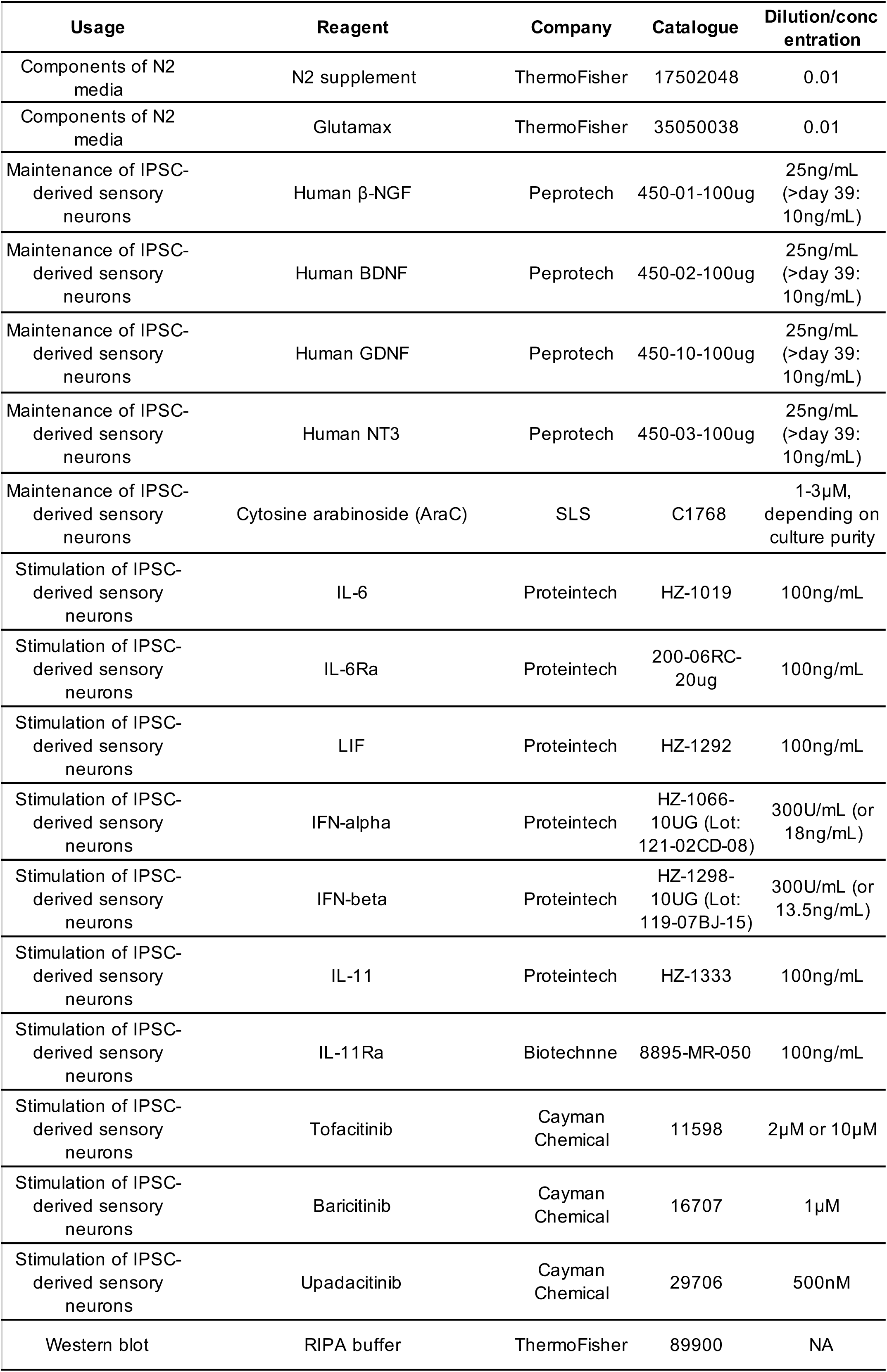
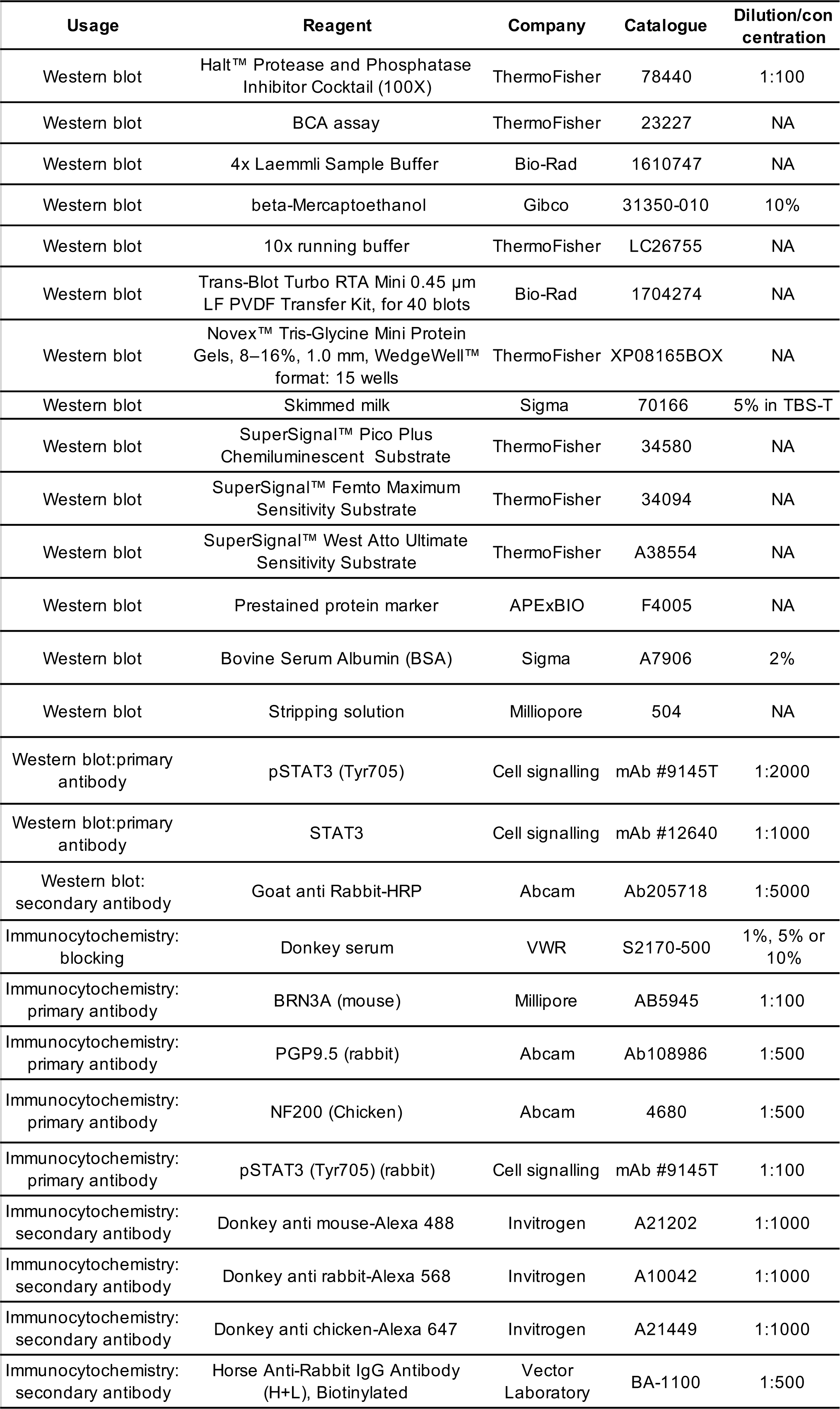
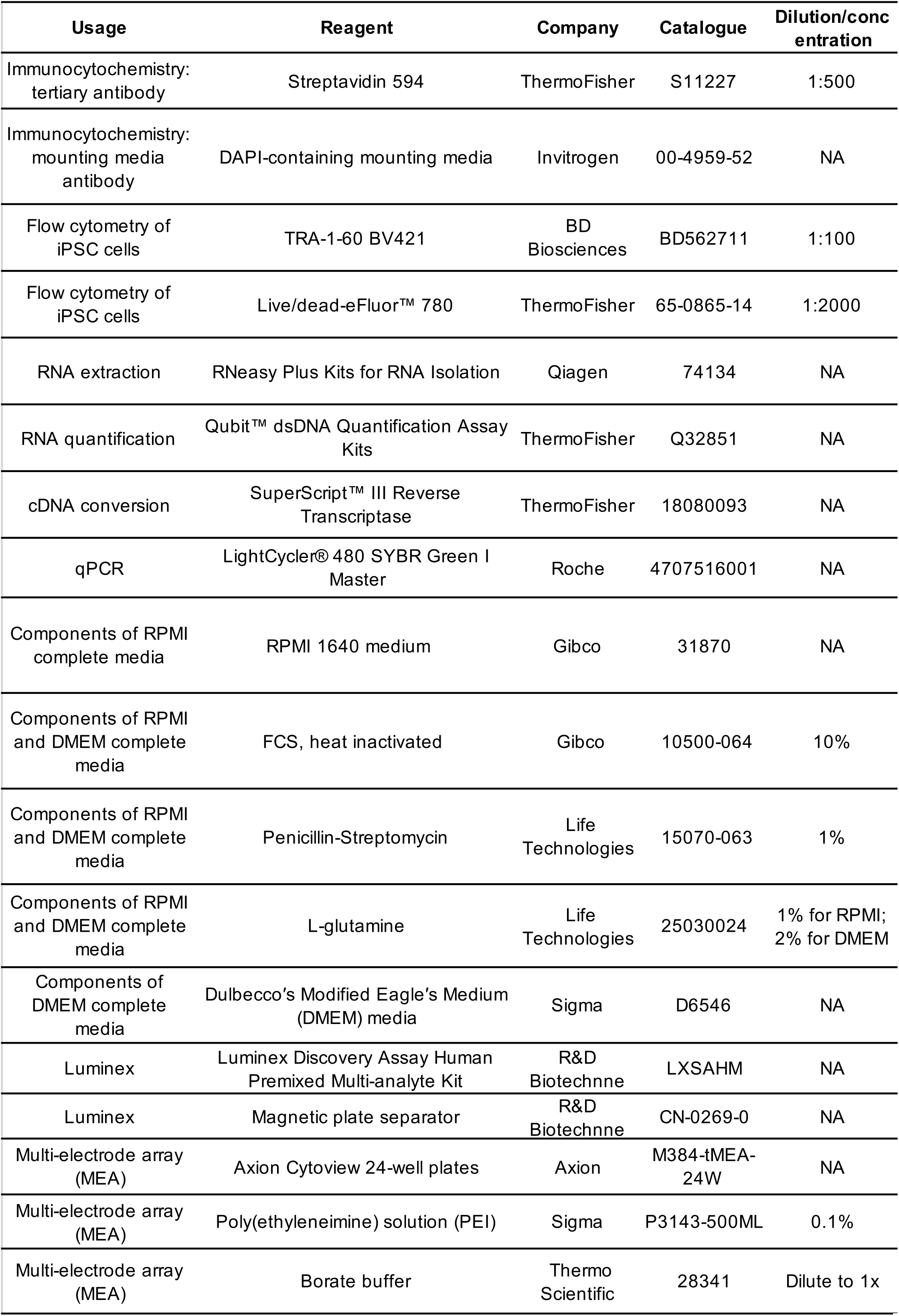
Reagents used in the study.

| Usage | Reagent | Company | Catalogue | Dilution/concentration |
| --- | --- | --- | --- | --- |
| IPSC maintenance | Stemflex media | ThermoFisher | A3349401 | NA |
| IPSC maintenance | Vitronectin | Stem cell technology | 7180 | 40µL vitronectin in 1mL PBS |
| IPSC maintenance | Geltrex | ThermoFisher | A1569601 | 1:100 for IPSC, 1:50 for neurons |
| IPSC maintenance | Versene | ThermoFisher | 15040066 | NA |
| IPSC maintenance and replating | TrypLE | ThermoFisher | 12604013 | NA |
| Replating | Rho Kinase inhibitor Y-27632 | Enzo | ALX-270-333-M005 | 10µM |
| Differentiation of IPSC into sensory neurons | Mouse Embryonic Fibroblast (MEF) Conditioned Media | Bio-techne | AR005 | NA |
| Differentiation of IPSC into sensory neurons | Human FGF-2 | Miltenyi | 130-093-839 | 10 ng/mL |
| Differentiation of IPSC into sensory neurons | SB431542 | Bio-techne | 1614-10 | 10µM |
| Differentiation of IPSC into sensory neurons | LDN-193189 | Bio-techne | 6053-10 | 100nM |
| Differentiation of IPSC into sensory neurons | CHIR99021 | Bio-techne | 4423-10 | 3µM |
| Differentiation of IPSC into sensory neurons | SU5402 | Merck | SML0443-5mg | 10µM |
| Differentiation of IPSC into sensory neurons | DAPT | Bio-techne | 2634-10 | 10µM |
| Components of Knockout DMEM media | Knockout DMEM basal | ThermoFisher | 10829018 | NA |
| Components of Knockout DMEM media | Knockout serum replacement | ThermoFisher | 10829018 | 15% |
| Components of Knockout DMEM media | Glutamax | ThermoFisher | 35050038 | 1% |
| Components of Knockout DMEM media | Nonessential amino acids | ThermoFisher | 11140035 | 1% |
| Components of Knockout DMEM media | Beta-Mercaptoethanol | Gibco | 31350-010 | 100 µM |
| Components of Knockout DMEM, N2 and stemflex media | Antibiotic/antimycotic | ThermoFisher | 15240096 | 1% |
| Components of N2 media | Neurobasal medium | ThermoFisher | 21103049 | NA |
| Components of N2 media | B27 supplement | ThermoFisher | 17504044 | 0.02 |

| Supplementary Table 1. Reagents used in the study (continued) |  |  |  |  |
| --- | --- | --- | --- | --- |
| Usage | Reagent | Company | Catalogue | Dilution/conc<br>entration |
| Components of N2 media | N2 supplement | ThermoFisher | 17502048 | 0.01 |
| Components of N2 media | Glutamax | ThermoFisher | 35050038 | 0.01 |
| Maintenance of IPSC-derived sensory neurons | Human β-NGF | Peprotech | 450-01-100ug | 25ng/mL (>day 39: 10ng/mL) |
| Maintenance of IPSC-derived sensory neurons | Human BDNF | Peprotech | 450-02-100ug | 25ng/mL (>day 39: 10ng/mL) |
| Maintenance of IPSC-derived sensory neurons | Human GDNF | Peprotech | 450-10-100ug | 25ng/mL (>day 39: 10ng/mL) |
| Maintenance of IPSC-derived sensory neurons | Human NT3 | Peprotech | 450-03-100ug | 25ng/mL (>day 39: 10ng/mL) |
| Maintenance of IPSC-derived sensory neurons | Cytosine arabinoside (AraC) | SLS | C1768 | 1-3μM, depending on culture purity |
| Stimulation of IPSC-derived sensory neurons | IL-6 | Proteintech | HZ-1019 | 100ng/mL |
| Stimulation of IPSC-derived sensory neurons | IL-6Ra | Proteintech | 200-06RC-20ug | 100ng/mL |
| Stimulation of IPSC-derived sensory neurons | LIF | Proteintech | HZ-1292 | 100ng/mL |
| Stimulation of IPSC-derived sensory neurons | IFN-alpha | Proteintech | HZ-1066-10UG (Lot: 121-02CD-08) | 300U/mL (or 18ng/mL) |
| Stimulation of IPSC-derived sensory neurons | IFN-beta | Proteintech | HZ-1298-10UG (Lot: 119-07BJ-15) | 300U/mL (or 13.5ng/mL) |
| Stimulation of IPSC-derived sensory neurons | IL-11 | Proteintech | HZ-1333 | 100ng/mL |
| Stimulation of IPSC-derived sensory neurons | IL-11Ra | Biotechnne | 8895-MR-050 | 100ng/mL |
| Stimulation of IPSC-derived sensory neurons | Tofacitinib | Cayman Chemical | 11598 | 2μM or 10μM |
| Stimulation of IPSC-derived sensory neurons | Baricitinib | Cayman Chemical | 16707 | 1μM |
| Stimulation of IPSC-derived sensory neurons | Upadacitinib | Cayman Chemical | 29706 | 500nM |
| Western blot | RIPA buffer | ThermoFisher | 89900 | NA |
| Usage | Reagent | Company | Catalogue | Dilution/concentration |
| Western blot | Halt™ Protease and Phosphatase Inhibitor Cocktail (100X) | ThermoFisher | 78440 | 1:100 |
| Western blot | BCA assay | ThermoFisher | 23227 | NA |
| Western blot | 4x Laemmli Sample Buffer | Bio-Rad | 1610747 | NA |
| Western blot | beta-Mercaptoethanol | Gibco | 31350-010 | 10% |
| Western blot | 10x running buffer | ThermoFisher | LC26755 | NA |
| Western blot | Trans-Blot Turbo RTA Mini 0.45 µm LF PVDF Transfer Kit, for 40 blots | Bio-Rad | 1704274 | NA |
| Western blot | Novex™ Tris-Glycine Mini Protein Gels, 8–16%, 1.0 mm, WedgeWell™ format: 15 wells | ThermoFisher | XP08165BOX | NA |
| Western blot | Skimmed milk | Sigma | 70166 | 5% in TBS-T |
| Western blot | SuperSignal™ Pico Plus Chemiluminescent Substrate | ThermoFisher | 34580 | NA |
| Western blot | SuperSignal™ Femto Maximum Sensitivity Substrate | ThermoFisher | 34094 | NA |
| Western blot | SuperSignal™ West Atto Ultimate Sensitivity Substrate | ThermoFisher | A38554 | NA |
| Western blot | Prestained protein marker | APExBIO | F4005 | NA |
| Western blot | Bovine Serum Albumin (BSA) | Sigma | A7906 | 2% |
| Western blot | Stripping solution | Milliopore | 504 | NA |
| Western blot:primary antibody | pSTAT3 (Tyr705) | Cell signalling | mAb #9145T | 1:2000 |
| Western blot:primary antibody | STAT3 | Cell signalling | mAb #12640 | 1:1000 |
| Western blot:secondary antibody | Goat anti Rabbit-HRP | Abcam | Ab205718 | 1:5000 |
| Immunocytochemistry: blocking | Donkey serum | VWR | S2170-500 | 1%, 5% or 10% |
| Immunocytochemistry: primary antibody | BRN3A (mouse) | Millipore | AB5945 | 1:100 |
| Immunocytochemistry: primary antibody | PGP9.5 (rabbit) | Abcam | Ab108986 | 1:500 |
| Immunocytochemistry: primary antibody | NF200 (Chicken) | Abcam | 4680 | 1:500 |
| Immunocytochemistry: primary antibody | pSTAT3 (Tyr705) (rabbit) | Cell signalling | mAb #9145T | 1:100 |
| Immunocytochemistry: secondary antibody | Donkey anti mouse-Alexa 488 | Invitrogen | A21202 | 1:1000 |
| Immunocytochemistry: secondary antibody | Donkey anti rabbit-Alexa 568 | Invitrogen | A10042 | 1:1000 |
| Immunocytochemistry: secondary antibody | Donkey anti chicken-Alexa 647 | Invitrogen | A21449 | 1:1000 |
| Immunocytochemistry: secondary antibody | Horse Anti-Rabbit IgG Antibody (H+L), Biotinylated | Vector Laboratory | BA-1100 | 1:500 |
| Usage | Reagent | Company | Catalogue | Dilution/conc<br>entration |
| Immunocytochemistry:<br>tertiary antibody | Streptavidin 594 | ThermoFisher | S11227 | 1:500 |
| Immunocytochemistry:<br>mounting media<br>antibody | DAPI-containing mounting media | Invitrogen | 00-4959-52 | NA |
| Flow cytometry of<br>iPSC cells | TRA-1-60 BV421 | BD<br>Biosciences | BD562711 | 1:100 |
| Flow cytometry of<br>iPSC cells | Live/dead-eFluor™ 780 | ThermoFisher | 65-0865-14 | 1:2000 |
| RNA extraction | RNeasy Plus Kits for RNA Isolation | Qiagen | 74134 | NA |
| RNA quantification | Qubit™ dsDNA Quantification Assay<br>Kits | ThermoFisher | Q32851 | NA |
| cDNA conversion | SuperScript™ III Reverse<br>Transcriptase | ThermoFisher | 18080093 | NA |
| qPCR | LightCycler® 480 SYBR Green I<br>Master | Roche | 4707516001 | NA |
| Components of RPMI<br>complete media | RPMI 1640 medium | Gibco | 31870 | NA |
| Components of RPMI<br>and DMEM complete<br>media | FCS, heat inactivated | Gibco | 10500-064 | 10% |
| Components of RPMI<br>and DMEM complete<br>media | Penicillin-Streptomycin | Life<br>Technologies | 15070-063 | 1% |
| Components of RPMI<br>and DMEM complete<br>media | L-glutamine | Life<br>Technologies | 25030024 | 1% for RPMI;<br>2% for DMEM |
| Components of<br>DMEM complete<br>media | Dulbecco's Modified Eagle's Medium<br>(DMEM) media | Sigma | D6546 | NA |
| Luminex | Luminex Discovery Assay Human<br>Premixed Multi-analyte Kit | R&D<br>Biotechnne | LXSAHM | NA |
| Luminex | Magnetic plate separator | R&D<br>Biotechnne | CN-0269-0 | NA |
| Multi-electrode array<br>(MEA) | Axion Cytoview 24-well plates | Axion | M384-tMEA-24W | NA |
| Multi-electrode array<br>(MEA) | Poly(ethyleneimine) solution (PEI) | Sigma | P3143-500ML | 0.1% |
| Multi-electrode array<br>(MEA) | Borate buffer | Thermo<br>Scientific | 28341 | Dilute to 1x |

**Supplementary Table 2.**
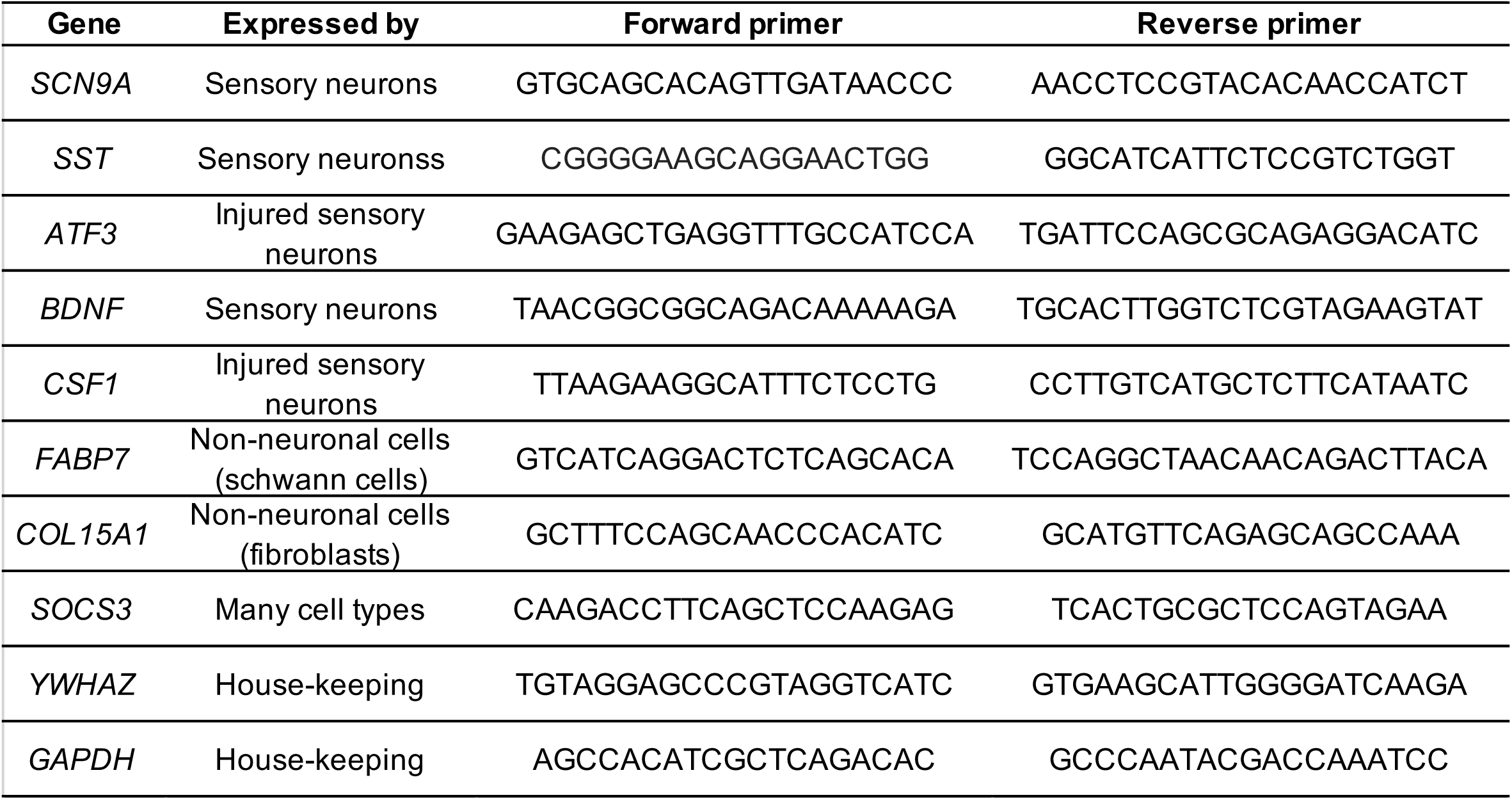
Primers used in this study. All were tested for their efficiency and specificity, and the correct product size was confirmed via gel electrophoresis.

**Supplementary Table 3.** Statistical tests performed in this study, including unit of analysis and observed effect size.

| Targets | Comparison | Unit of analysis | n | Observed effect size (d) | Statistical test | p-value | p-value cut-off after multiple comparison correction | Significant (*) |
| --- | --- | --- | --- | --- | --- | --- | --- | --- |
| pSTAT3 | SF vs SF+tofa | Culture well: Kute4 JK T2 | 6 | 1.2 | Paired nonparametric t-test | 0.01256 | 0.05 | * |
| IL-6 | RA serum vs RA SF | Patient sample | 12 | 0.98 | Paired nonparametric t-test | 0.00049 | 0.008 | * |
| LIF | RA serum vs RA SF | Patient sample | 12 | 1.08 | Paired nonparametric t-test | 0.00244 | 0.008 | * |
| IL-11 | RA serum vs RA SF | Patient sample | 12 | 2.67 | Paired nonparametric t-test | 0.00049 | 0.008 | * |
| OSM | RA serum vs RA SF | Patient sample | 12 | -0.32 | Paired nonparametric t-test | 0.46973 | 0.008 | NS |
| IFN-alpha | RA serum vs RA SF | Patient sample | 12 | 1.75 | Paired nonparametric t-test | 0.00592 | 0.008 | * |
| IFN-beta | RA serum vs RA SF | Patient sample | 12 | 1.66 | Paired nonparametric t-test | 0.00592 | 0.008 | * |
| IL-6 | levels of mediator in RA SF vs neuronal pSTAT3/STAT3 | Patient sample | 10 | 0.79 | Spearman's correlation | 0.003 | 0.008 | * |
| LIF | levels of mediator in RA SF vs neuronal pSTAT3/STAT3 | Patient sample | 10 | 0.82 | Spearman's correlation | 0.002 | 0.008 | * |
| IL-11 | levels of mediator in RA SF vs neuronal pSTAT3/STAT3 | Patient sample | 10 | 0.45 | Spearman's correlation | 0.09 | 0.008 | NS |
| OSM | levels of mediator in RA SF vs neuronal pSTAT3/STAT3 | Patient sample | 10 | 0.43 | Spearman's correlation | 0.11 | 0.008 | NS |
| IFN-alpha | levels of mediator in RA SF vs neuronal pSTAT3/STAT3 | Patient sample | 10 | 0.8 | Spearman's correlation | 0.003 | 0.008 | * |
| IFN-beta | levels of mediator in RA SF vs neuronal pSTAT3/STAT3 | Patient sample | 10 | 0.15 | Spearman's correlation | 0.34 | 0.008 | NS |
| SOCS3 | IL-6+sIL-6R vs Ctrl | Culture well: Kute4 T35 UKB T9, T10 | 9 | 6.17 | One-tailed nonparametric t-test | <0.0001 | 0.0125 | * |
| ATF3 | IL-6+sIL-6R vs Ctrl | Culture well: Kute4 T35 UKB T9, T10 | 9 | 1.13 | One-tailed nonparametric t-test | 0.0094 | 0.0125 | * |
| BDNF | IL-6+sIL-6R vs Ctrl | Culture well: Kute4 T35 UKB T9, T10 | 9 | 0.91 | One-tailed nonparametric t-test | 0.017 | 0.0125 | NS |
| CSF1 | IL-6+sIL-6R vs Ctrl | Culture well: Kute4 T35 UKB T9, T10 | 9 | 2.61 | One-tailed nonparametric t-test | 0.0001 | 0.0125 | * |
| SOCS3 | IL-6+sIL-6R vs IL-6+sIL-6R+tofa | Culture well: Kute4 T40, UKB T12 | 6 | 3.17 | One-tailed nonparametric t-test | 0.001 | 0.0125 | * |
| ATF3 | IL-6+sIL-6R vs IL-6+sIL-6R+tofa | Culture well: Kute4 T40, UKB T12 | 6 | 1.10 | One-tailed nonparametric t-test | 0.09 | 0.0125 | NS |
| BDNF | IL-6+sIL-6R vs IL-6+sIL-6R+tofa | Culture well: Kute4 T40, UKB T12 | 6 | 0.65 | One-tailed nonparametric t-test | 0.002 | 0.0125 | * |
| CSF1 | IL-6+sIL-6R vs IL-6+sIL-6R+tofa | Culture well: Kute4 T40, UKB T12 | 6 | 2.62 | One-tailed nonparametric t-test | 0.004 | 0.0125 | * |
| SOCS3 | LIF vs LIF+tofa | Culture well: Kute4 T40, UKB T12 | 6 | 2.59 | One-tailed nonparametric t-test | 0.001 | 0.0125 | * |
| ATF3 | LIF vs LIF+tofa | Culture well: Kute4 T40, UKB T12 | 6 | 1.78 | One-tailed nonparametric t-test | 0.001 | 0.0125 | * |
| BDNF | LIF vs LIF+tofa | Culture well: Kute4 T40, UKB T12 | 6 | 1.29 | One-tailed nonparametric t-test | 0.001 | 0.0125 | * |
| CSF1 | LIF vs LIF+tofa | Culture well: Kute4 T40, UKB T12 | 6 | 1.56 | One-tailed nonparametric t-test | 0.02 | 0.0125 | NS |
| Mean firing rate | LIF & temperature as independent variables | Culture well: Kute4 T40, UKB T14 | 14-16 | NA | Repeated measures ANOVA | LIF (0.045), temperature(0.067) | NA | * |

Observed effect size was calculated as Cohen's $d = (M_2 - M_1) / \sqrt{((SD_1^2 + SD_2^2) / 2)}$ . In experiments where multiple statistical tests were performed, Bonferroni correction was performed. When the observed p value was lower than the new p-value cut-off, \* was added. NS: Not significant. Wilcoxon signed-rank tests were used for paired nonparametric t tests and Mann Whitney U test for the one-tailed, independent nonparametric t tests.

**Supplementary Table 4.** Demographic (and clinical) parameters of RA.

| Parameters | RA | HC |
| --- | --- | --- |
| Sex (F/M) | 12/0 | 4/0 |
| Age (years):<br>mean(SD) | 59(19) | 37.5(13) |
| Disease duration (years):<br>mean(SD) | 12.3(10.4) | NA |
| ACPA: +/- | 6/0 | NA |
| Rheumatoid factor: +/- | 12/0 | NA |
| Erosive: +/- | 8/1 | NA |
| DAS28:<br>mean(SD) | 5.1(1.3) | NA |
| TJC: mean(SD) | 6.1(4.8) | NA |
| SJC: mean(SD) | 5.6(5.2) | NA |
| ESR: mean(SD) | 34.6(24) | NA |
| CRP: mean(SD) | 16.8(12.6) | NA |

**ACPA:** Anti-citrullinated protein antibodies (ACPA). **DAS28:** Disease Activity Score 28. **TJC:** tender joint counts. **SJC:** swollen joint counts. **ESR:** Erythrocyte sedimentation rate (ESR) (mm/hr). **CRP:** C-reactive protein (mg/L). Number of individuals with missing information: Age (1), Erosive (3), TJC (4), SJC(4), ESR(4), CRP(1), ACPA(6).

**Supplementary Table 5.** Luminex assay information.

|  | HC serum<br>(pg/mL):<br>Mean<br>(SD) | RA serum<br>(pg/mL):<br>Mean<br>(SD) | RA SF<br>(pg/mL):<br>Mean<br>(SD) | Standard1<br>(pg/mL) | Standard6<br>(pg/mL) | Sensitivity<br>(pg/mL) | Standard<br>curve-fit<br>R^2 |
| --- | --- | --- | --- | --- | --- | --- | --- |
| IL-6 | 130.0<br>(112.0) | 179.4<br>(382.0) | 17734.4<br>(25368.1) | 79000 | 325.1 | 170.0 | 0.997 |
| IFN-beta | 1.9<br>(0.0) | 1.9<br>(0.0) | 21.0<br>(16.1) | 6880 | 28.3 | 1.0 | 0.903 |
| IL-6R<br>alpha | 36477.9<br>(5306.9) | 36286.4<br>(11374.5) | 36909.2<br>(12379.4) | 116380 | 478.9 | 1.2 | 0.998 |
| Oncostatin M/OSM | 7952.3<br>(581.6) | 7740.9<br>(839.4) | 7324.8<br>(1653.3) | 186460 | 767.3 | 88.6 | 0.962 |
| LIF | 12.2<br>(8.8) | 15.1<br>(10.1) | 57.1<br>(54.3) | 11780 | 48.5 | 18.6 | 0.996 |
| IFN-alpha | 1.3<br>(0.0) | 1.9<br>(2.2) | 62.0<br>(48.5) | 5400 | 22.2 | 0.5 | 0.999 |
| IL-11 | 182.1<br>(0.0) | 478.3<br>(707.1) | 3310.5<br>(1324.0) | 213600 | 879 | 49.4 | 0.926 |

HC serum, RA serum and RA SF were diluted 1:2 and 1:100 and run on the same Luminex plate. 1:100 dilution wells were chosen to calculate IL-6 concentration and 1:2 wells were chosen to calculate the concentration for the rest of the analytes, based on which dilution had most datapoints fall within the standard curve (STD1-STD6). The standard curve was derived using a five parameter logistic (5-PL) curve-fit, calculated by xPonent software upon running the plate on a FLEXMAP 3D machine according to manufacturer's instructions. Standard 1 (top standard), Standard 6 (bottom standard) and sensitivity have taken the corresponding dilution factors into account. Sensitivity information was obtained on the R&D website when designing the panel: [biotechne.com//rl/t9MRQ6Nj](https://www.rndsystems.com/products/assays/assay-panels.aspx).

## Supplementary methods

### 1. Bioinformatic analyses

We derived information from five previously published RNA sequencing datasets: two single-nuclei sequencing of post-mortem human sensory neurons, GSE168243 and GSE201586 (1, 2), two bulk RNA sequencing datasets of IPSC-derived sensory neurons, one of which was of neurons we grew in-house (GSE268585) and a compilation dataset (3). For the single-nuclei RNA sequencing datasets, we generated pseudobulk profiles for each biological replicate and plotted the expression of our genes of interest.

### 2. Fibroblast and PBMC/SFMC culture

Cryopreserved RA synovial fibroblasts were thawed and cultured at 37°C with 5% CO2 in DMEM complete medium, containing 10% FCS, 1% penicillin/streptomycin, 2% L-glutamine and 1μg/mL Amphotericin B/Fungizone. Three RA patient-derived fibroblast lines at passage 5-6 were used in this study. Fibroblasts were seeded in 12-well plates (60,000 cells/well) the night before the experiment and subsequently stimulated with or without IL-1beta for 24 hours.

Cryopreserved healthy donor and RA patient PBMC and/or SFMC were thawed and cultured in 24-well plates (1 million cells/well) in RPMI complete medium only or with plate-bound anti-CD3/soluble anti-CD28 mAbs for 3 days at 37°C with 5% CO2. Complete RPMI medium was composed of RPMI 1640 medium containing 10% FCS, 1% Penicillin/Streptomycin and 1% L-glutamine. Conditioned media were harvested by spinning down the cells at 5900g for 3.5 minutes. Supernatants were removed and stored at −80°C until Luminex assessment.

### 3. Sensitivity analyses

For Luminex experiments, using n=12 patient samples, we conducted paired non-parametric t-tests to compare the levels of our target cytokines in RA serum and SF. Since we measured six cytokines, the p-value cut-off was lowered to 0.008 using Bonferroni correction. Sensitivity analysis suggests this design will provide an 80% chance to detect effects size of d=1.26 or larger. Using n=10 patient samples, we also correlated the levels of mediators in SF with the levels of neuronal pSTAT3/STAT3 they induced as measured by Western blot. p=0.008 was used as a cut-off, with sensitivity analysis indicating that this will provide an 80% chance of seeing correlations of rho=0.83 or larger.

For gene expression analyses, using n=6-9 culture wells, we conducted one-tailed non-parametric t-tests to see whether IL-6+sIL-6R or LIF increased the expression of *SOCS3*, *ATF3*, *BDNF* and *CSF1*. Since four genes were tested, the p-value cut-off was lowered to 0.0125 using Bonferroni correction. Sensitivity analyses suggest this design has an 80% chance to detect effects of d=1.62-2.11.

