## Supplementary material for "Blockade of rheumatoid arthritis synovial fluid-induced sensory neuron activation by JAK inhibitors": Western blots raw files

### All Western blots used in manuscript

Yuening Li

2024.8

Note: Only pSTAT3 and STAT3 blots were used in the manuscript.
The ECL kit to image the blots are indicated on the lower right corners.
The relevant lanes used in the manuscript are indicated in each slide and marked by:

**Figure 3A**

Used in manuscript: RA1, ….., RA10, No stim

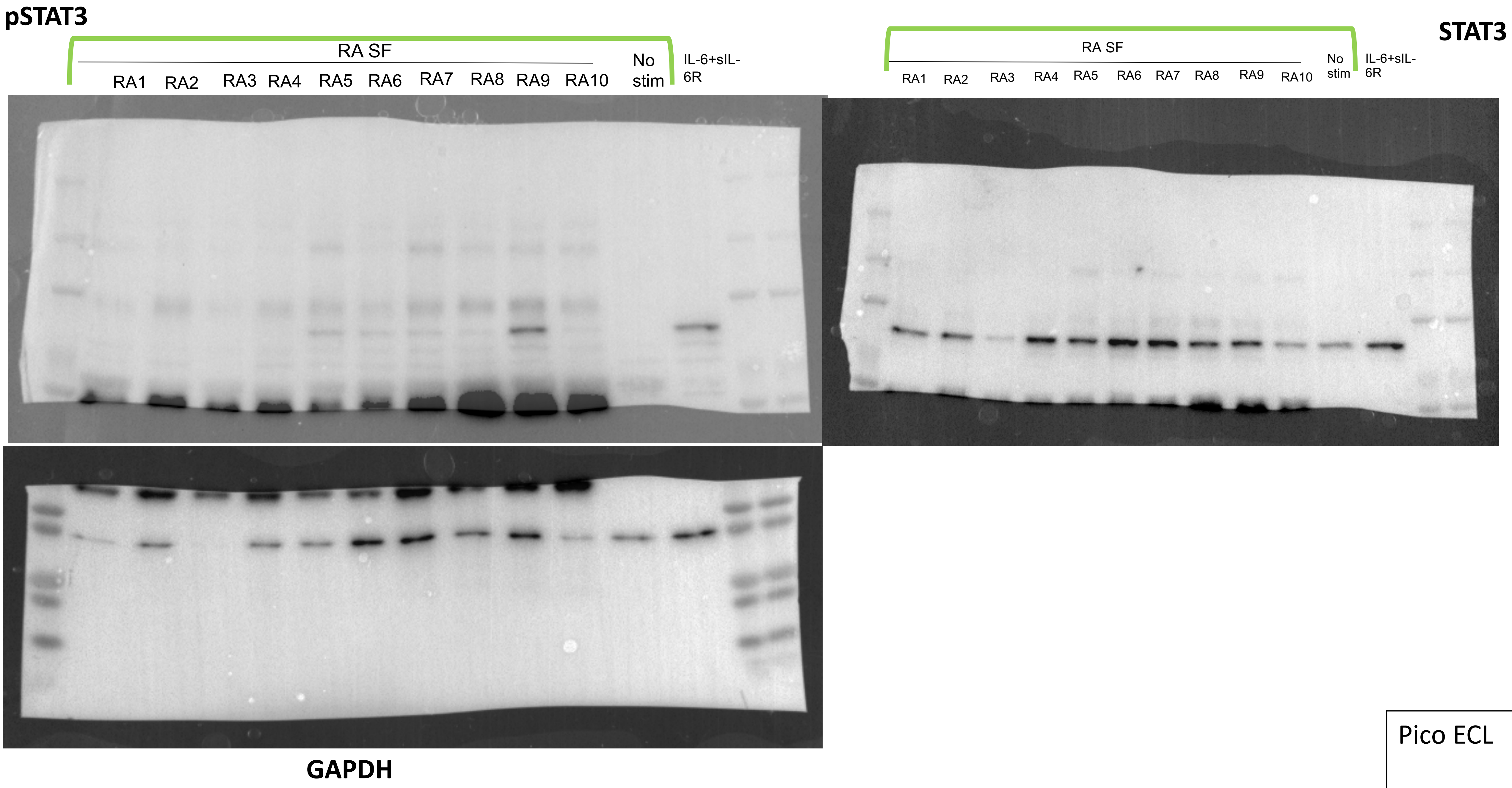


Pico ECL

**Figure 3B**

Used in manuscript: RA10, RA10 tofa, RA11, RA11 tofa, RA8, RA8, tofa

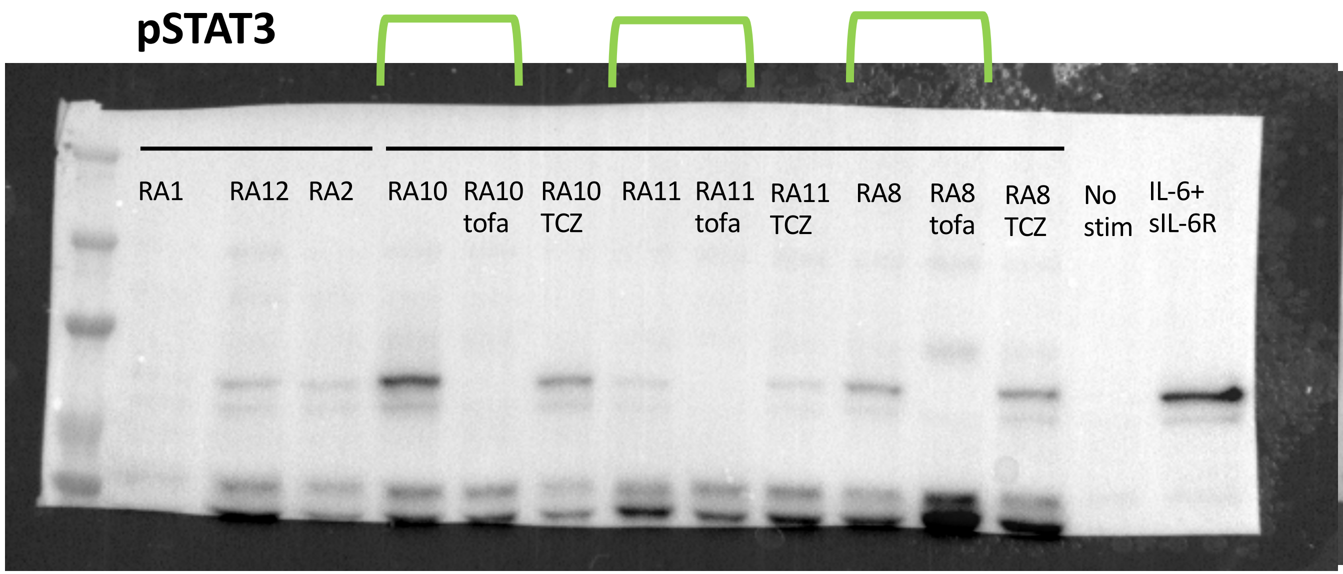


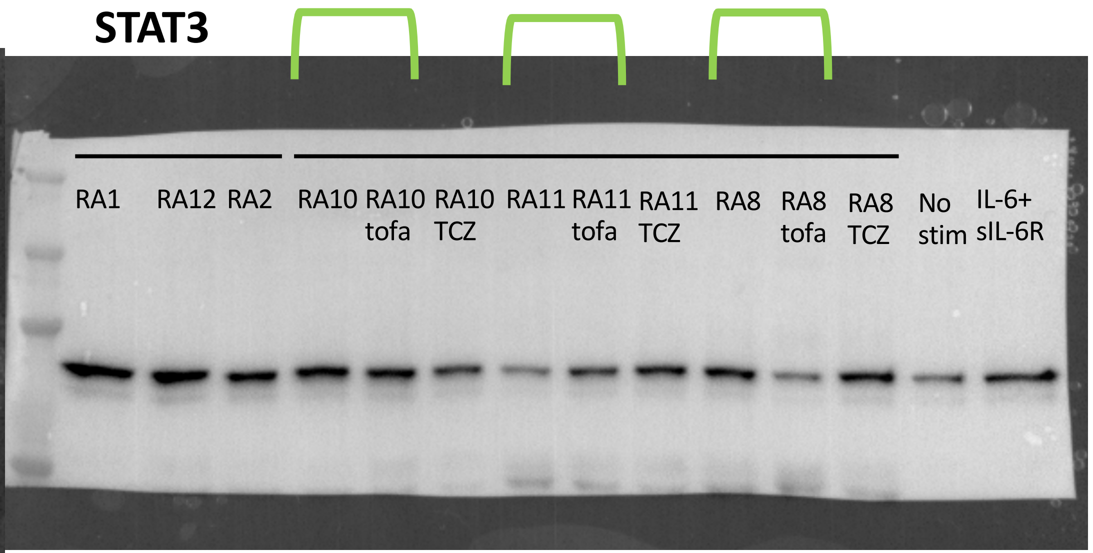


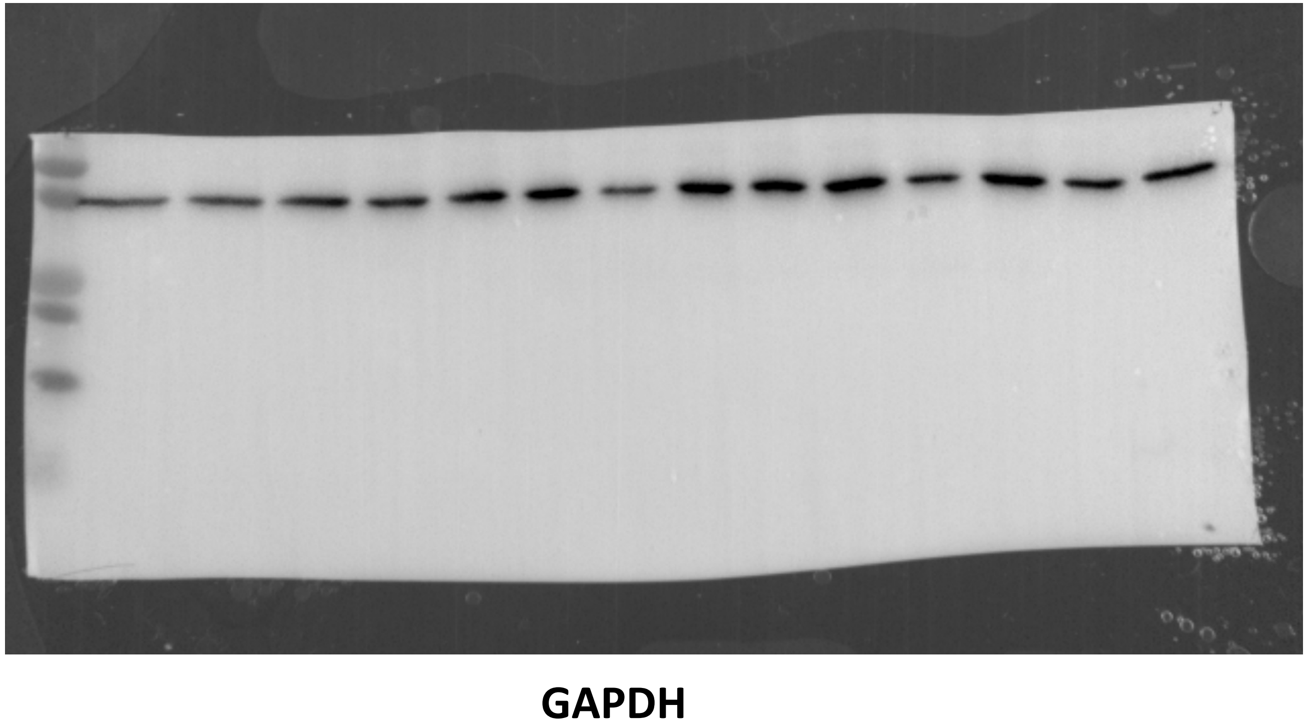


Tofa: tofacitinib
TCZ: tocilizumab

Pico ECL

### Figure 5A

Used in manuscript: all conditions

**pSTAT3**

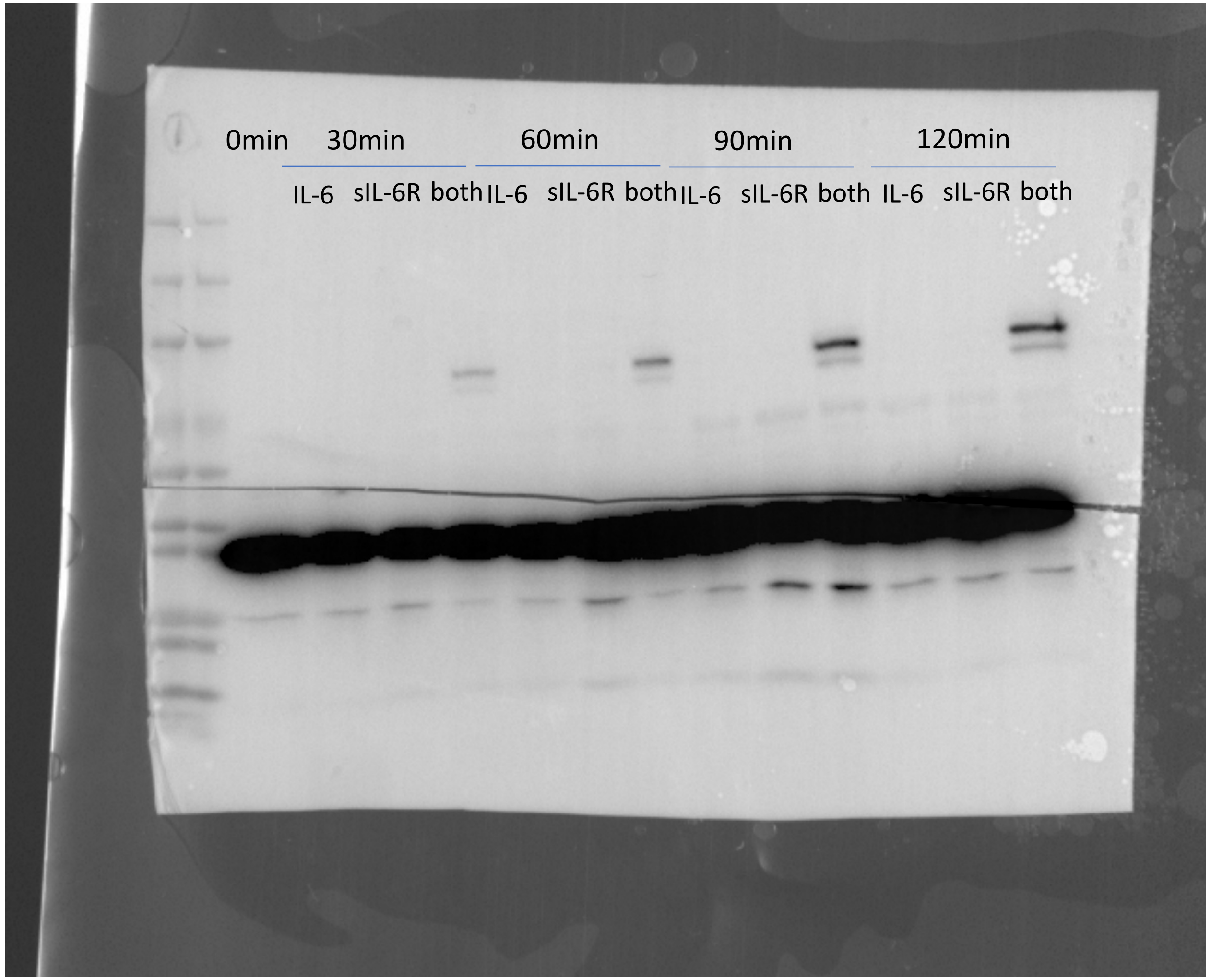


**GAPDH**

**STAT3**

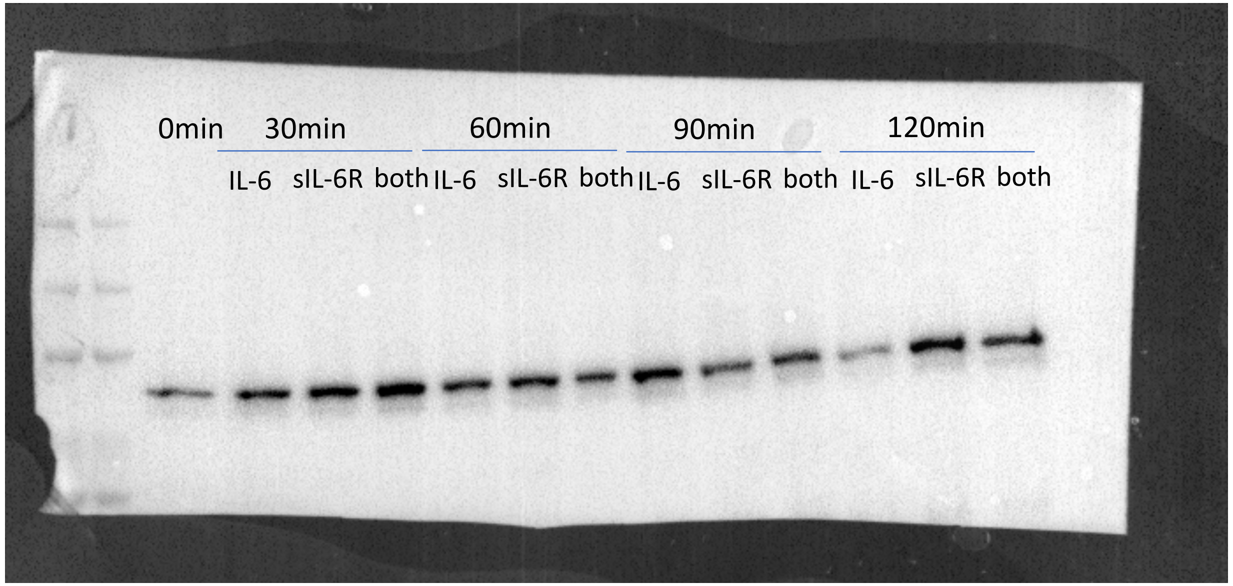


Pico ECL

**Figure 5B**

Used in manuscript: LIF lanes

**pSTAT3**

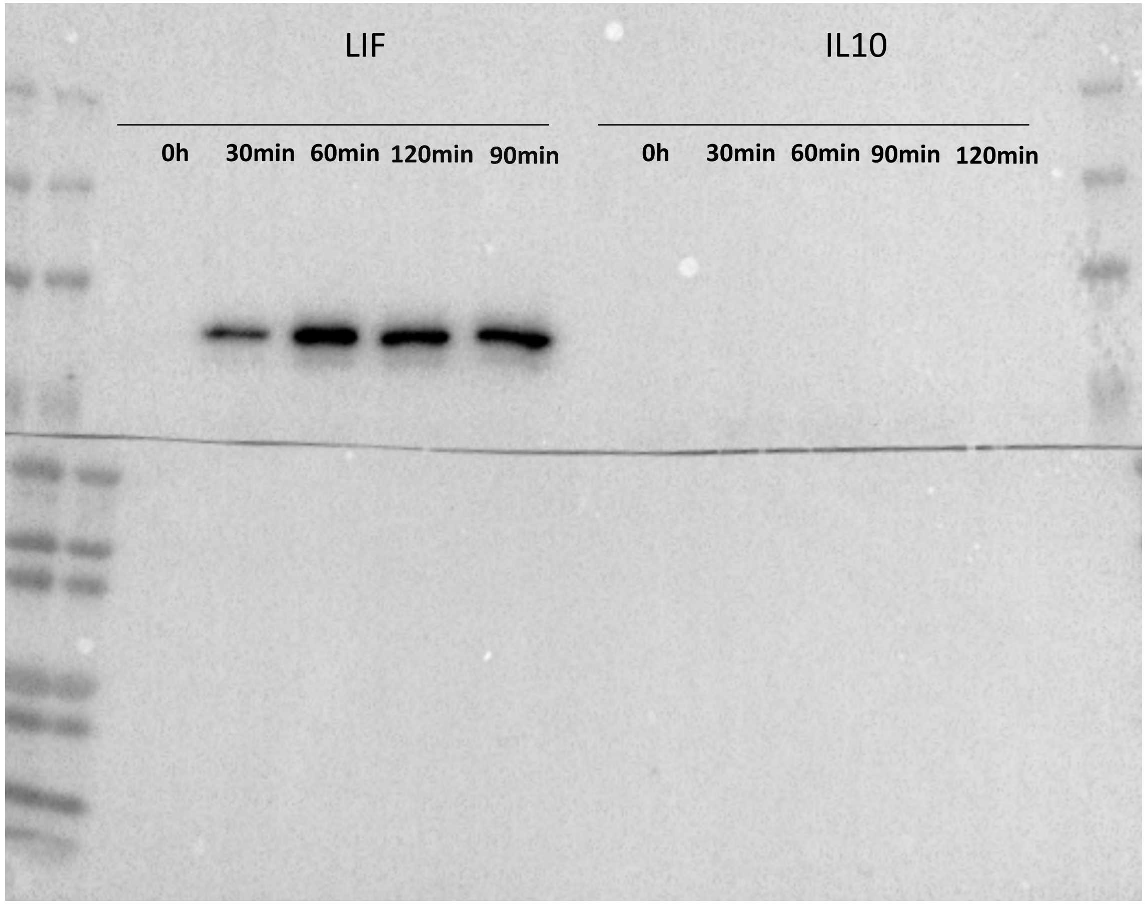


**STAT3**

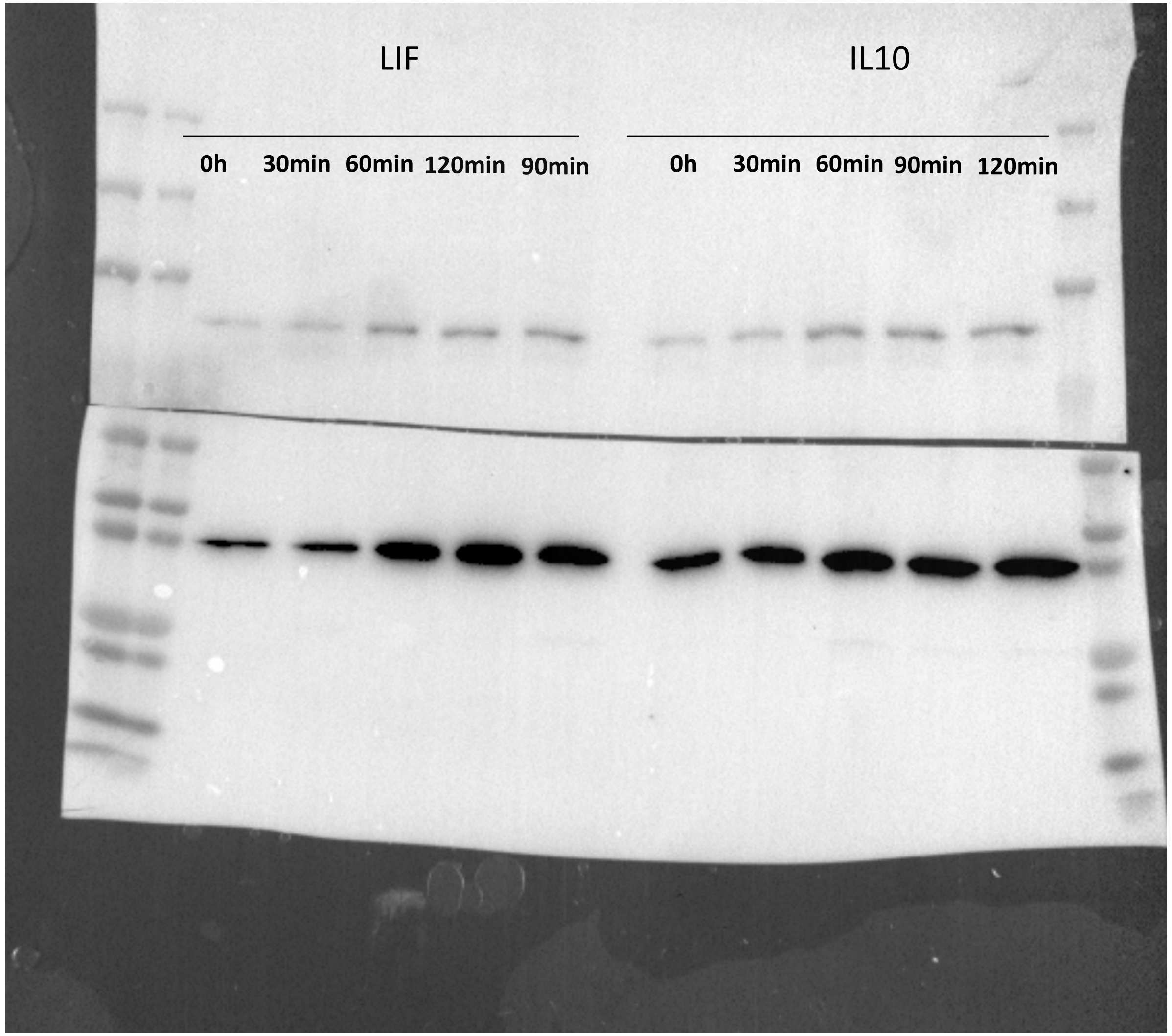


**GAPDH**

Pico ECL

**Figure 5C**

Used in manuscript: IFNα & IFNβ conditions

**pSTAT3**

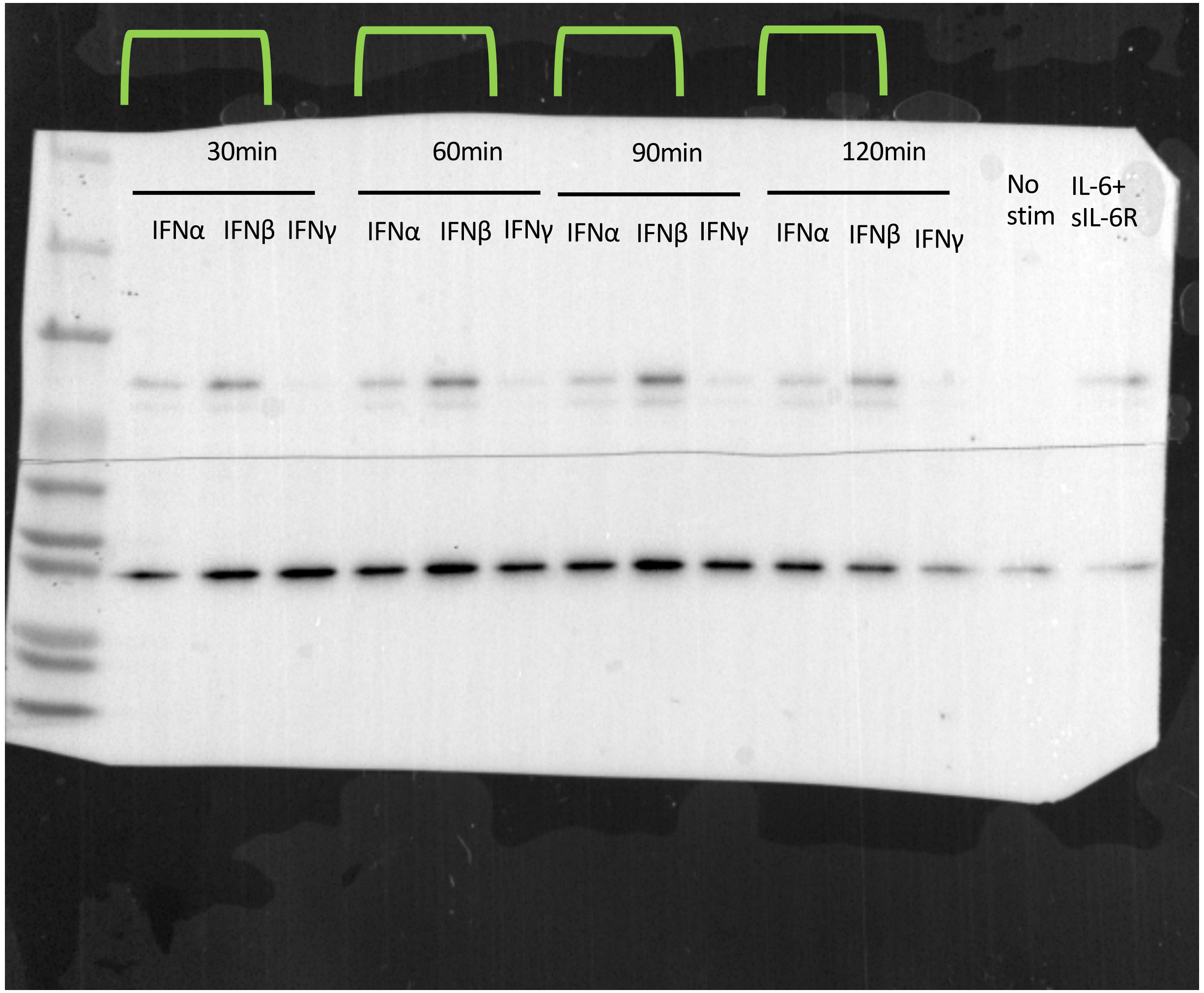


**GAPDH**

**STAT3**

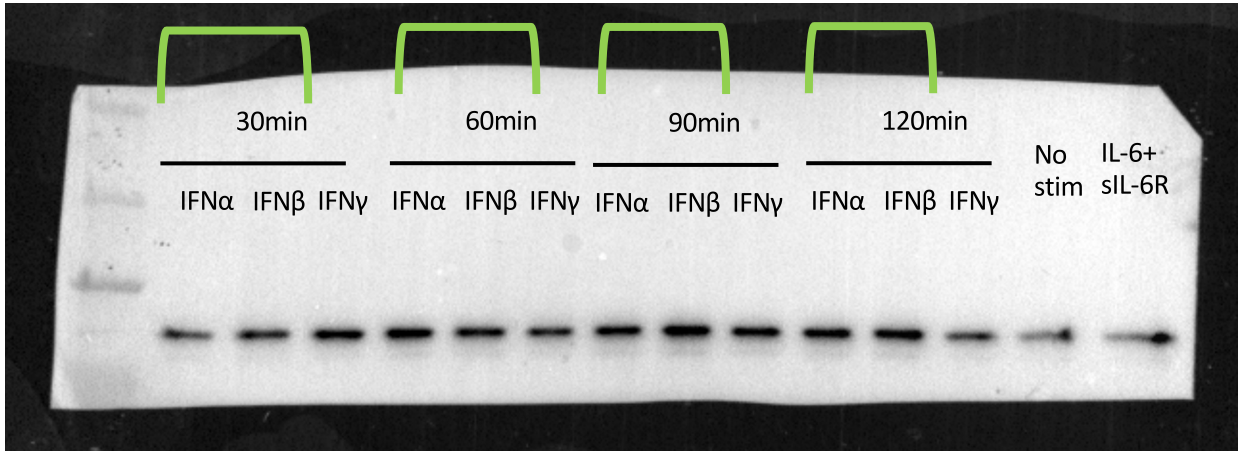


pSTAT3: Atto ECL
STAT3: Femto ECL

**Figure 5D**

Used in manuscript: 30min to 120min conditions

pSTAT3

STAT3

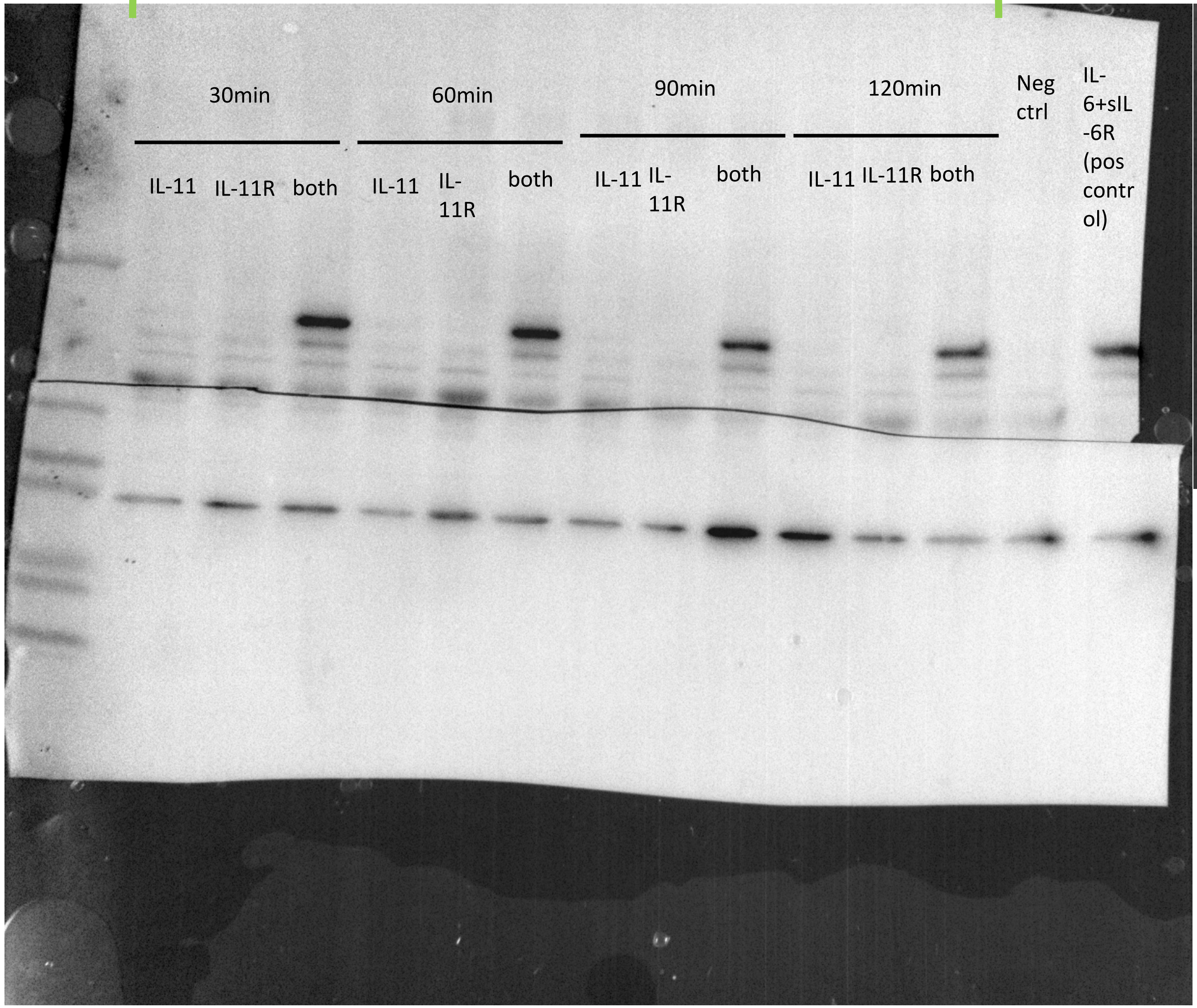


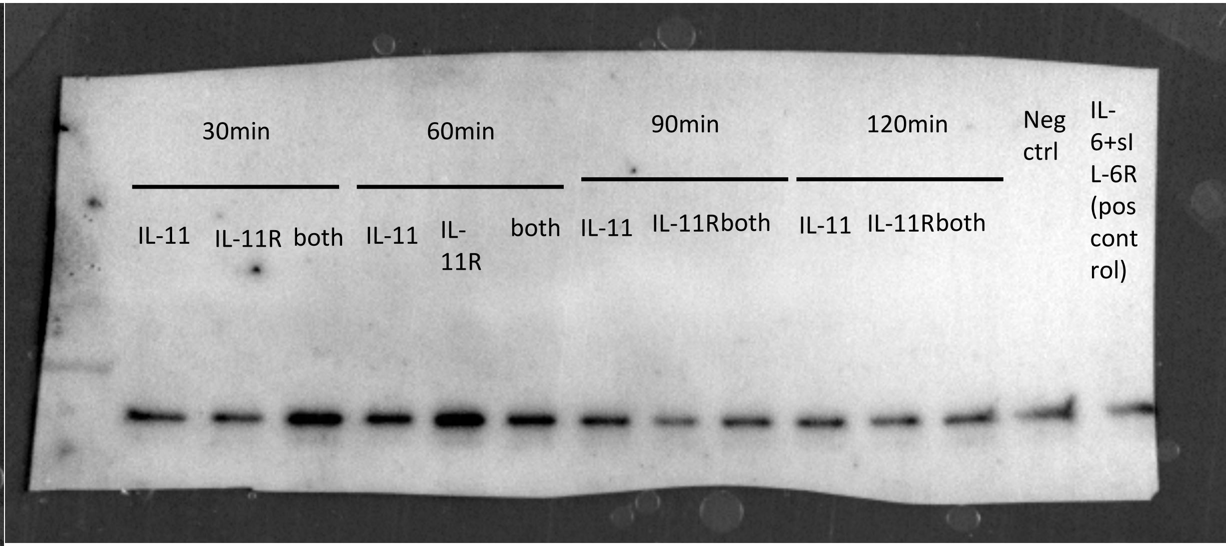


GAPDH

pSTAT3: Atto ECL
STAT3: Femto ECL

**Figure 5E**

Used in manuscript: No stim, IL-6+sIL-6R, LIF conditions

**pSTAT3**

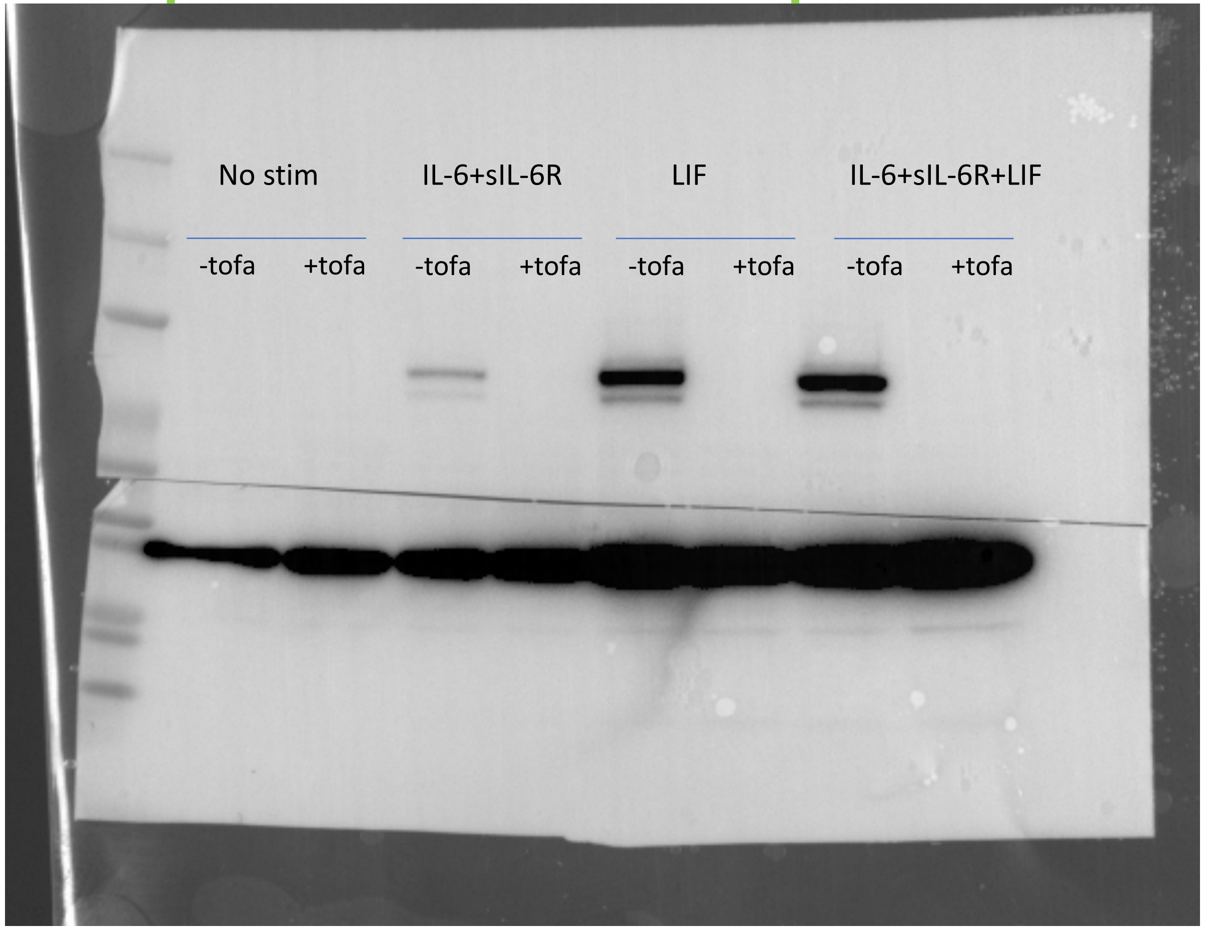


**GAPDH**

**STAT3**

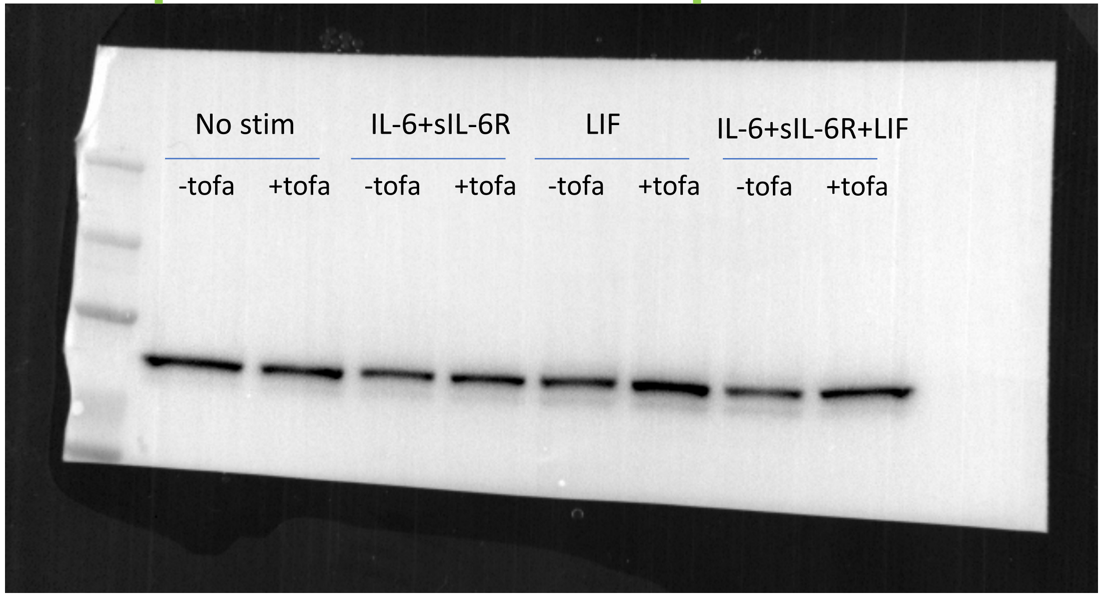


Pico ECL

**Supplementary Fig. 2A**

Used in manuscript: all but IL-6+sIL-6R condition

**pSTAT3**

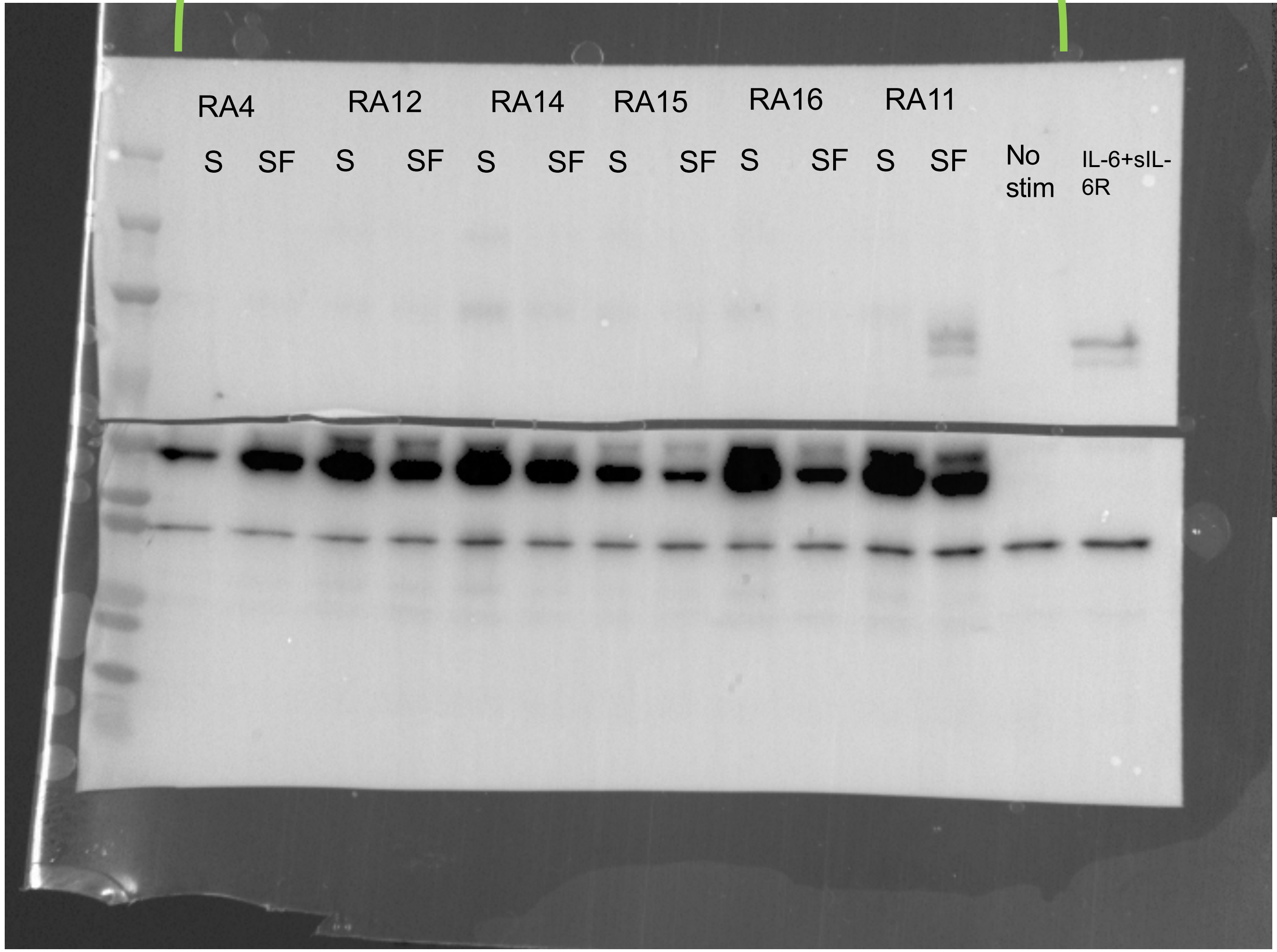


**STAT3**

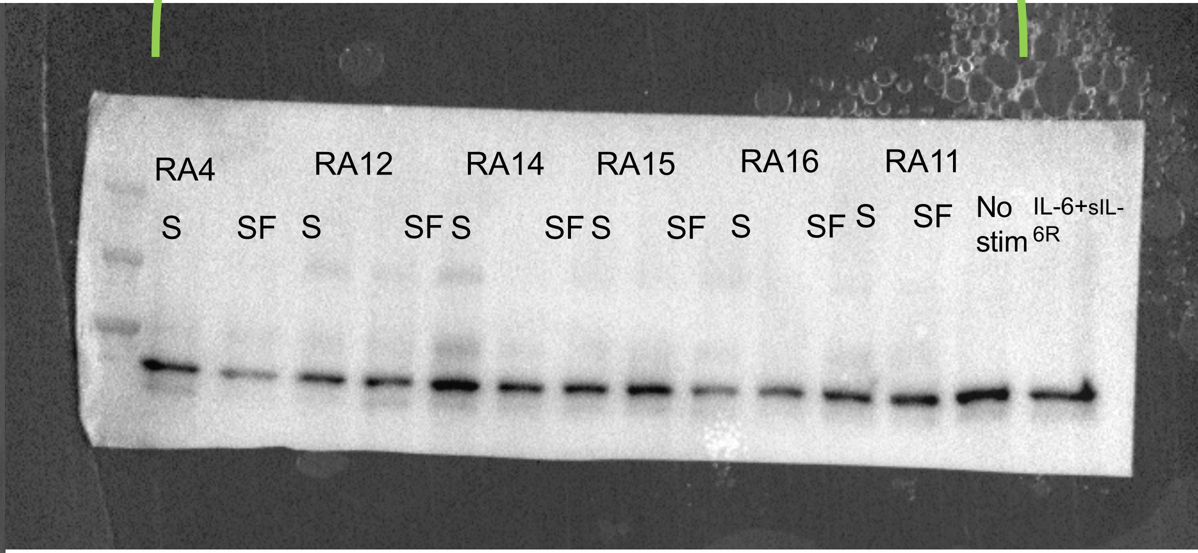


**GAPDH**

S: serum

Pico ECL

**Supplementary Figure 6A**

Used in manuscript: all but the last two lanes

**pSTAT3**

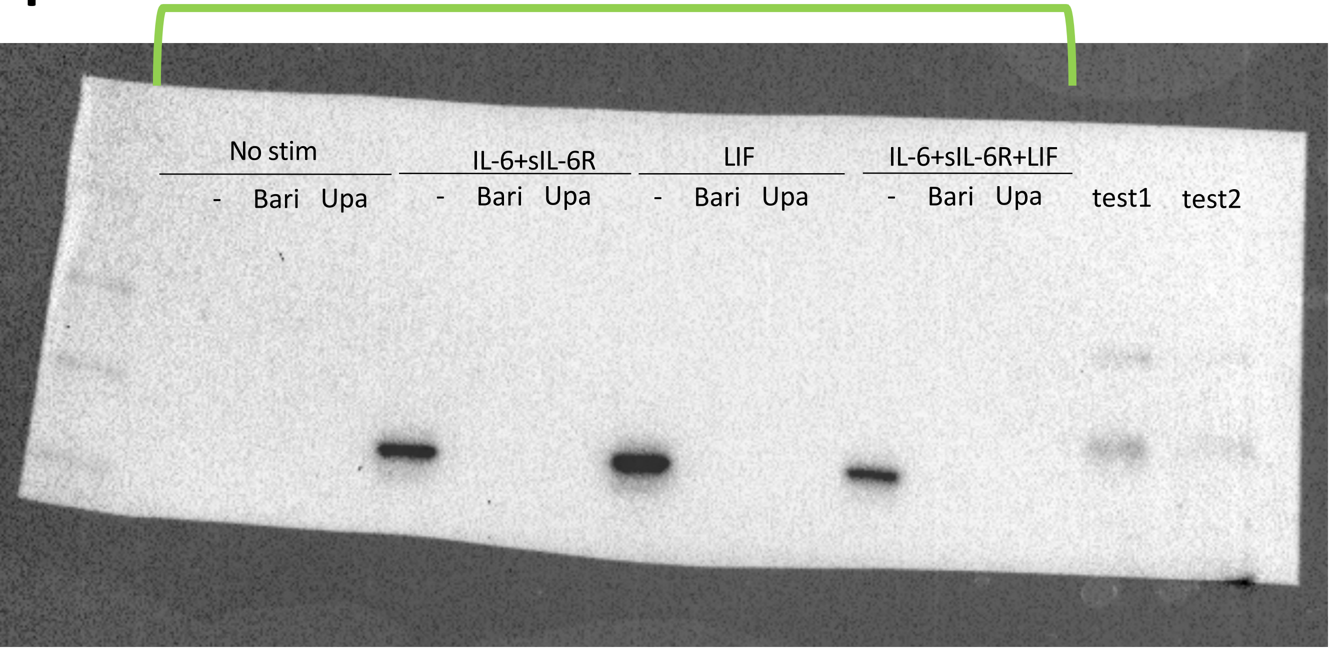


**STAT3**

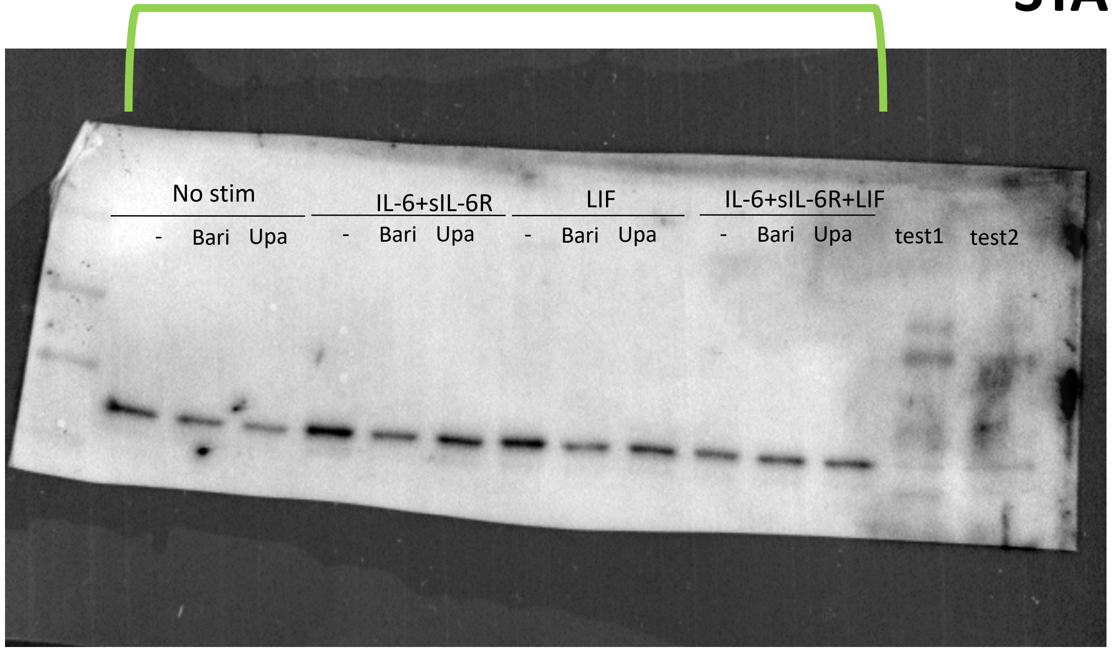


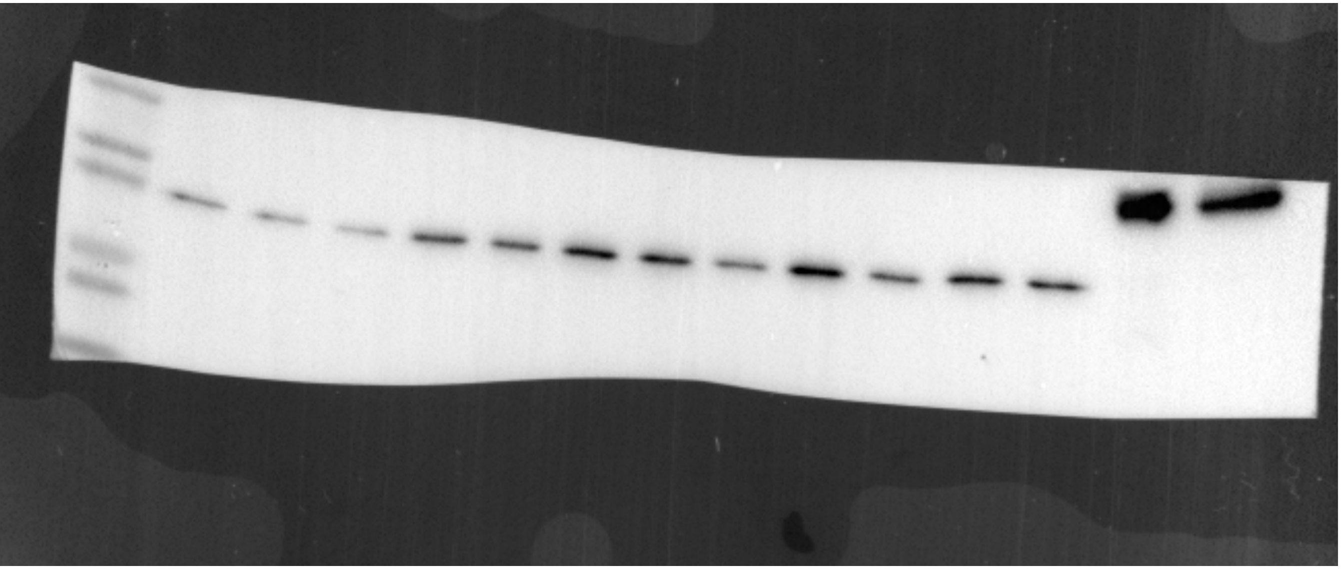

**GAPDH**

pSTAT3: pico ECL
STAT3: atto ECL
